# mRNA editing of the Alzheimer’s risk gene *APOE*

**DOI:** 10.64898/2026.09.03.749263

**Authors:** Valérie Blanc, Samantha J. Griffiths, Jürgen Haas, Nicholas O. Davidson, Richard Lathe

## Abstract

Variants in the human *APOE* gene govern the risk of Alzheimer’s disease and other disorders. Three major *APOE* variants in humans reflect C→T replacements at two positions in a single exon: an upstream variant (AE4 site) that differs between the ancestral APOE *ε4* allele (*APOE4*) (C) and human-specific *APOE2*/*E3* (T), and a downstream variant (AE2 site) that differentiates *APOE3*/*E4*(C)from *APOE2* (T). It has long been assumed that *APOE* allelotypes are genomically encoded, but here we report that multiple individuals express brain *APOE* C or U/T variant transcripts that differ from genomically templated versions. We demonstrate up to 10% C→U or U→C nucleotide replacement at AE4 and AE2, but not at other sites, and with no corresponding changes in genomic DNA. Single-cell transcriptomic datasets from brain microglia revealed sporadic (up to ∼8%) C→U replacement at AE2. We found 0.4–1.6% of brain transcripts in human *APOE* knock-in mice harbor selective C→U changes at either AE4 or AE2 sites. Transfection of HepG2 or Huh7 cells with either mouse or human APOBEC1 led to efficient (>90%) C→U editing of *APOE4* mRNA at the AE4 but not AE2 site, with lower (<10%) C→T editing of genomic DNA at the AE4 site by mouse, but not human, APOBEC1. Furthermore, interrogation of proteomic datasets revealed up to 4% of non-genomically encoded APOE peptides in human plasma, indicating that the edited *APOE* transcripts are functional *in vivo*. These data suggest that *APOE* mRNA is subject to RNA editing that interconverts the different allelic forms of *APOE*.

**Author summary:** Human *APOE* gene variants govern the risk of Alzheimer’s disease (AD) and other disorders. Three alleles are widespread: ancestral *E4* and human-specific *E3* and *E2*. AD risk declines in the order *E4* > *E3* > *E2*. It has been assumed that the *APOE* allotype we inherit is laid down at birth, but we report that *APOE* mRNA is enzymatically edited to convert *E4* to *E3*/*E2*, and/or *E2* to *E3*/*E4*. Up to ∼10% conversion was seen in brain, and up to 100% *in vitro* driven by the RNA-editing enzyme APOBEC1. Proteomic analysis of human plasma argues that edited *APOE* transcripts are functional *in vivo*.

## Introduction

Apolipoprotein E (APOE) is a 34 kDa secreted and cell-associated immunomodulator and lipid transport protein that modulates the risk of Alzheimer’s disease (AD). The human *APOE* locus on chromosome 19 is highly polymorphic, and three allelic variants, *APOE4, E3*, and *E2*, are widespread in the population. Individuals who harbor two *APOE4* alleles are up to 10-fold more likely to develop AD than individuals with two *APOE2* alleles, and *E3* carriers are at intermediate risk (1, 2). There are two polymorphic sites in *APOE* – a upstream site (C/T at rs429358; Cys112 versus Arg112) that differentiates the APOE4 (Arg) protein from E2 and E3 (Cys), designated here the AE4 site, and a downstream site (C/T at rs7412) that encodes Cys158 (in APOE2 protein) versus Arg158 (in APOE3 and E4), designated the AE2 site. Theoretically there should be four alleles (Cys-Cys, Cys-Arg, Arg-Cys, and Arg-Arg), but the two sites are in proximity, and because of linkage disequilibrium *APOE2* + *E3* + *E4* represent close to 100% of all alleles in human populations.

In addition to modulating the risk of AD and several other disorders (3), as well as longevity (4), APOE plays an important role in innate immunity (5). The primary phenotype of *Apoe*-deficient mice is susceptibility to infection (e.g., (6)), and in human the *APOE4* allele increases the severity of multiple infectious diseases. AD is increasingly associated with infection/inflammation (7), and the antimicrobial properties of APOE (8-10) resemble those reported for AD amyloid β (Aβ) (11); APOE also binds tightly to Aβ, and APOE aggregates are found abundantly in AD plaques (12). Because of the central involvement of APOE in innate immunity, the hypothesis has been advanced that *APOE* is a classic example of balancing selection (13), whereby different alleles of genes, particularly those involved in host–pathogen interactions, are maintained in the population to ensure diversity in the face of varying environmental challenge (14).

In our analysis of the contribution of different *APOE* genotypes to AD risk we discovered that some individuals express allelic variants that differ from their underlying genotype. Our findings suggest that the specific sites that discriminate between the ancestral *APOE4* gene and the human-specific *E3*/*E2* alleles are also subject to RNA editing.

## Results

### Unexpected allelic mRNA distributions at the human *APOE* locus

The *APOE2* and *E3* alleles are recent acquisitions by the human lineage (Fig. 1A). To explore their contribution to AD, samples of four different regions of brain (amygdala, BA24, hippocampus, and hypothalamus) from controls and patients with AD from the Edinburgh Brain Bank (EBB) underwent RNA sequencing (RNA-seq; datasets reported in (15, 16)); samples of another region (cerebellum) from the same individuals were used for genomic DNA sequencing. For haplotype analysis, the RNA-seq datasets were examined using oligonucleotide query sequences (’probes’) that discriminate between *APOE4, E3*, and *E2* (Methods). We predicted that all transcripts would either (i) match the C or U/T versions (homozygosity: both alleles are identical) or (ii) manifest a 50/50 mixture (heterozygosity: two alleles are present at this site). Because RNA-seq data generate an output in which U in the source material is replaced by T, we use T to designate the identity of the nucleotide irrespective of whether the source was U or T, except in cases where the specific identity in RNA or DNA is relevant.

**Fig. 1.**
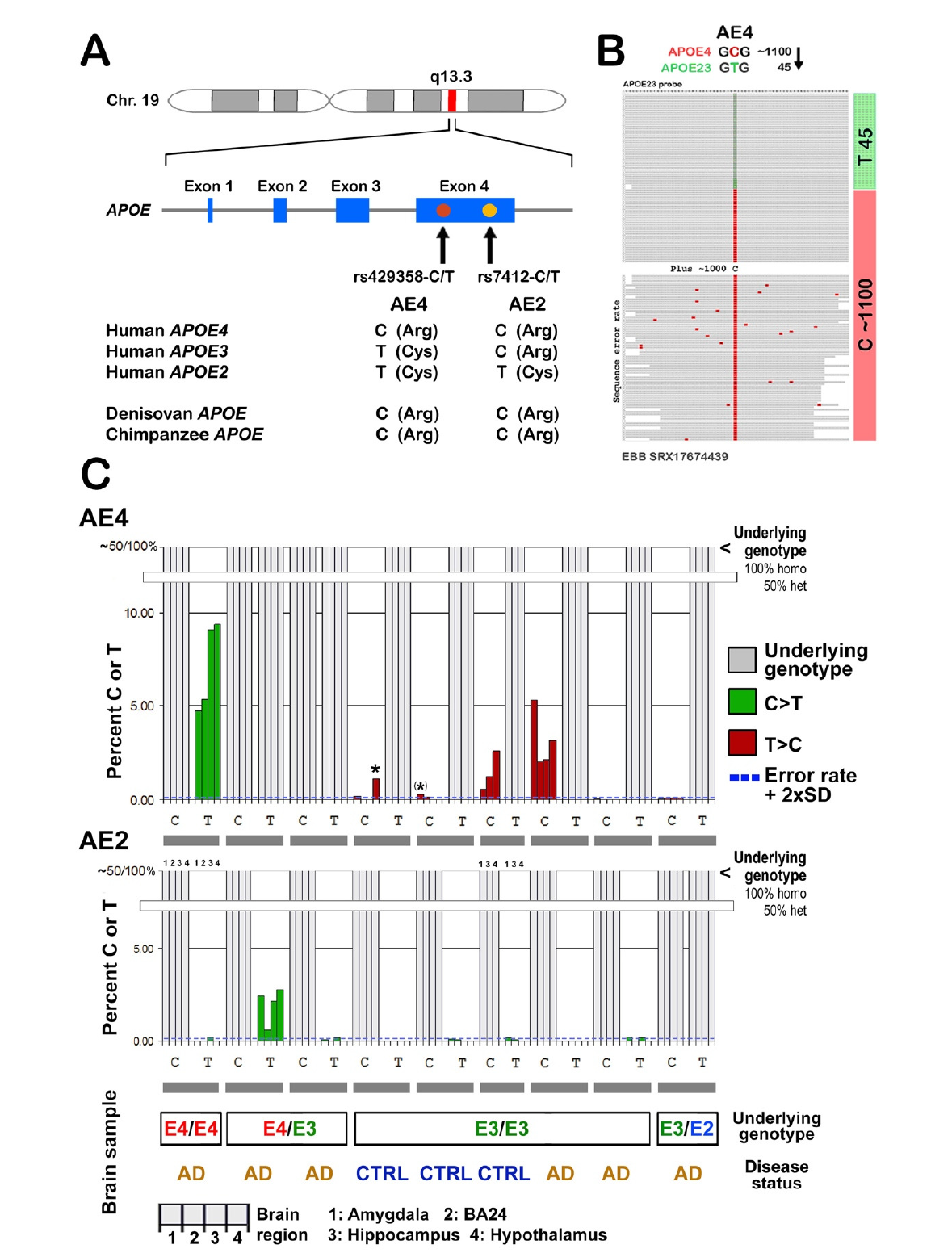
Human *APOE* gene and unexpected sequence variants in *APOE* mRNA in human brain. (A) Three major alleles of human *APOE*; panel inspired by (49). The two polymorphic sites in human *APOE* are designated here AE4 (distinguishes *APOE4* from *E2* + *E3*) and AE2 (distinguishes *APOE2* from *E3* + *E4*). *APOE* in other mammalian species including chimpanzee and the Denisovan hominin (50, 51) resembles human *APOE4*, suggesting that *E4* is the ancestral version whereas *E2* and *E3* are recent acquisitions by the human lineage. (B) BLAST searching of brain RNA-seq data with *APOE* probes (64-mers) revealed an unusually high frequency of T variants at the upstream polymorphic site, AE4, in an individual homozygous for the C variant (*APOE4*). Because this search employed a probe specific for *APOE2* + *E3* (’APO23’), BLAST displays the perfect matches first, followed by *APOE4*, short transcripts, and what are inferred to be sequencing errors. (C) Frequency of C→T (green) and T→C (red) changes versus the dominant genetic background at the AE4 and AE2 sites in *APOE* cDNA from brain from nine different individuals determined using allele-specific probes (36-mers). In two individuals only one sample gave evidence of sequence changes (indicated by asterisks), but both were significant because the frequency was above the sequence error rate + 2×SD (dotted blue line; i.e., *P* = <0.05). Samples (left to right) are amygdala, BA24, hippocampus, and hypothalamus, as indicated. Details of the individuals, samples, and datasets are given in Table S1. Abbreviations: AD, Alzheimer’s disease; CTRL, control without cognitive deficits; het, heterozygote; homo, homozygote; SD, standard deviation.

Our findings revealed divergence in *APOE* transcripts at both the AE4 and AE2 sites from the expected homozygous/heterozygous distribution, and long probes (64-mers) detected multiple T residues in brain cDNAs instead of the expected C residue (Fig. 1B). T→C changes were also observed in some samples (Fig. 1C). To refine the analysis, we used allele-specific sequences (36-mers) to probe brain cDNA, which revealed up to 9.36% C→T (green in Fig. 1C) and T→C (red) changes. Overall, the transcript sequences of six of nine individuals harbored variants that differed from the predicted genotype (Fig. 1C and Table S1). Of note, variant transcript distributions were individual-specific and were generally detected in multiple brain samples from the same individual, whereas brain samples from other individuals in the same series showed either no or limited evidence of sequence variants. In addition, there was no evidence for simultaneous changes at both AE4 and AE2 (Fig. 1C), although the underlying genotype may conceal some potential changes (e.g., C→T replacements at a particular site are not visible on either a heterozygous C/T or homozygous T/T genotype).

### Potential mechanisms for variation at the *APOE* locus

We first considered whether the results could be explained by cross-contamination or sequencing errors. However, up to 10% contamination would be necessary to explain the results, and genomic data (below) also argue against this possibility. To assess the potential role of sequencing errors, the distribution of variants in the *APOE* sequence was examined. The sequence differences in the AE4 region (64 nt) clustered specifically at the nucleotide that discriminates between *APOE4* and *E3*/*E2* (Fig. 2A). To confirm the specificity of the changes, 64-mer probes were designed for both the AE4 and AE2 regions in which each C or T pyrimidine nucleotide was systematically replaced by the other nucleotide (T or C), and the sequence datasets were rescreened. Except for a very small number of changes elsewhere, the changes occurred only at the AE4 and AE2 sites (Fig. 2B). Further evaluation of nucleotide-specific sequence changes in the AE4 region (Fig. 2C) revealed a mean error rate over the region of 0.047%, which is 70-fold below the observed frequency of single-nucleotide variants at the AE4 site seen in the EBB samples in Fig. 1 (mean 3.4%, range 0.6– 9.36%). The most common changes attributable to sequencing errors were A→C, and T→G (Fig. 2C), whereas the C→T changes seen in *APOE*, and particularly the T→C changes, were relatively rare. These findings suggest that sequencing errors are an unlikely source of the observed site-specific C↔T variations.

**Fig. 2.**
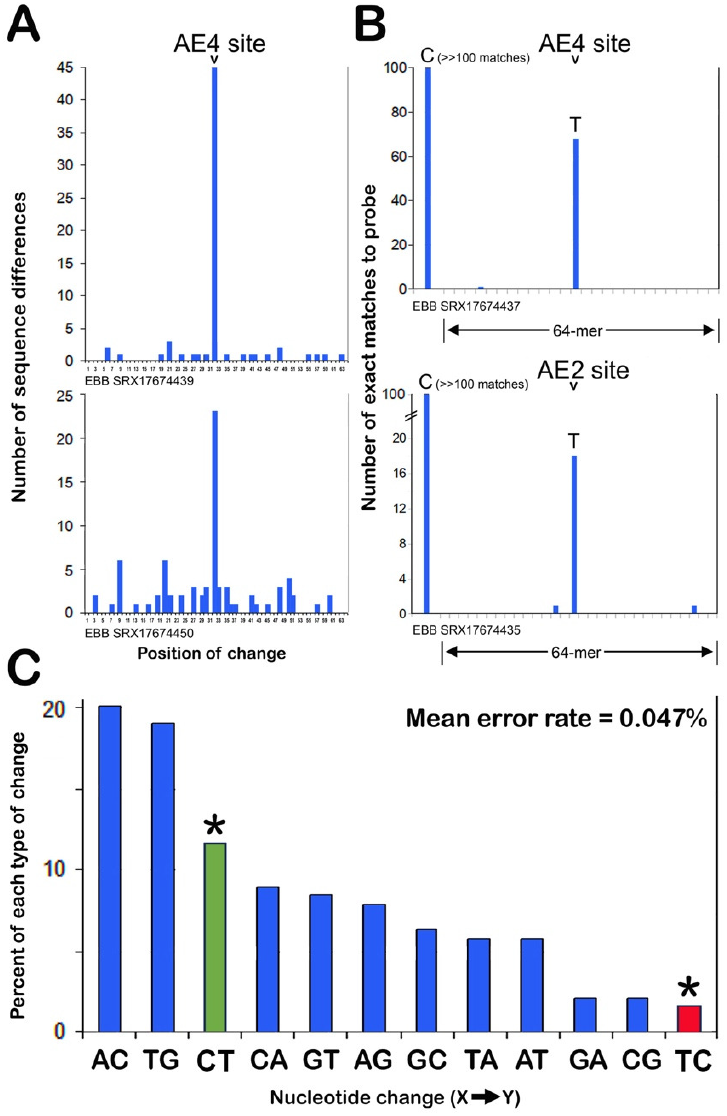
Location and type of sequence changes in *APOE*. (A) Plot of locations of all divergences from the canonical *APOE* sequence over a 64 nt region encompassing AE4, showing a high frequency of changes specifically at the AE4 site. (B) Experimental test of sequence errors: all C and T nucleotides in the 64-mer probes for AE4 and AE2 (Methods) were replaced by the other nucleotide (T or C), and BLAST searching was used to determine the frequency of each change. Excluding stochastic variants inferred to be sequencing errors (*N* = 3 in total), all pyrimidine/pyrimidine substitutions are at the specific sites in *APOE* that differentiate between the different alleles. (C) Types of sequence changes: excluding the AE4 site itself, the percentage and type of all divergence from the canonical sequence over the same region in all samples analyzed were calculated, showing that A→C and T→G changes are most frequent, C→T (green) is less abundant, and T→C changes (red) are the rarest. Edinburgh Brain Bank (EBB) sequence read archive (SRA/SRX) identities and corresponding biosamples are listed in Table S1.

### AE4 site changes are not present in genomic DNA

To address whether the observed changes are encoded in genomic DNA, specific primers were used to PCR amplify *APOE* exon 4 from brain DNA from the five individuals in Fig. 1 who manifested sequence changes. Two approaches were used. In the first, the sequencing chromatogram of each sample was quantified for C↔T changes. After subtraction of background, the proportion of T (or C) nucleotides in each lane (−1.5% to +1.08%), was calculated to be −0.61% (i.e., 0%) in an individual showing up to 10% C→T changes at the RNA level, demonstrating that the sequence changes are not present in genomic DNA (Fig. 3A). In a second approach >1000 individual DNA molecules were sequenced from the purified PCR products. We observed that variant sequences were systematically <1% of the expected C (or T) nucleotide (mean 0.24%) in genomic DNA whereas changes in RNA-derived cDNA were an order of magnitude more frequent (Fig. 3B,C). These findings support the conclusion that the sequence changes in the *APOE* transcript are not encoded in the genome and further suggest that cross-contamination of tissue samples is unlikely, because both RNA and DNA would be affected.

**Fig. 3.**
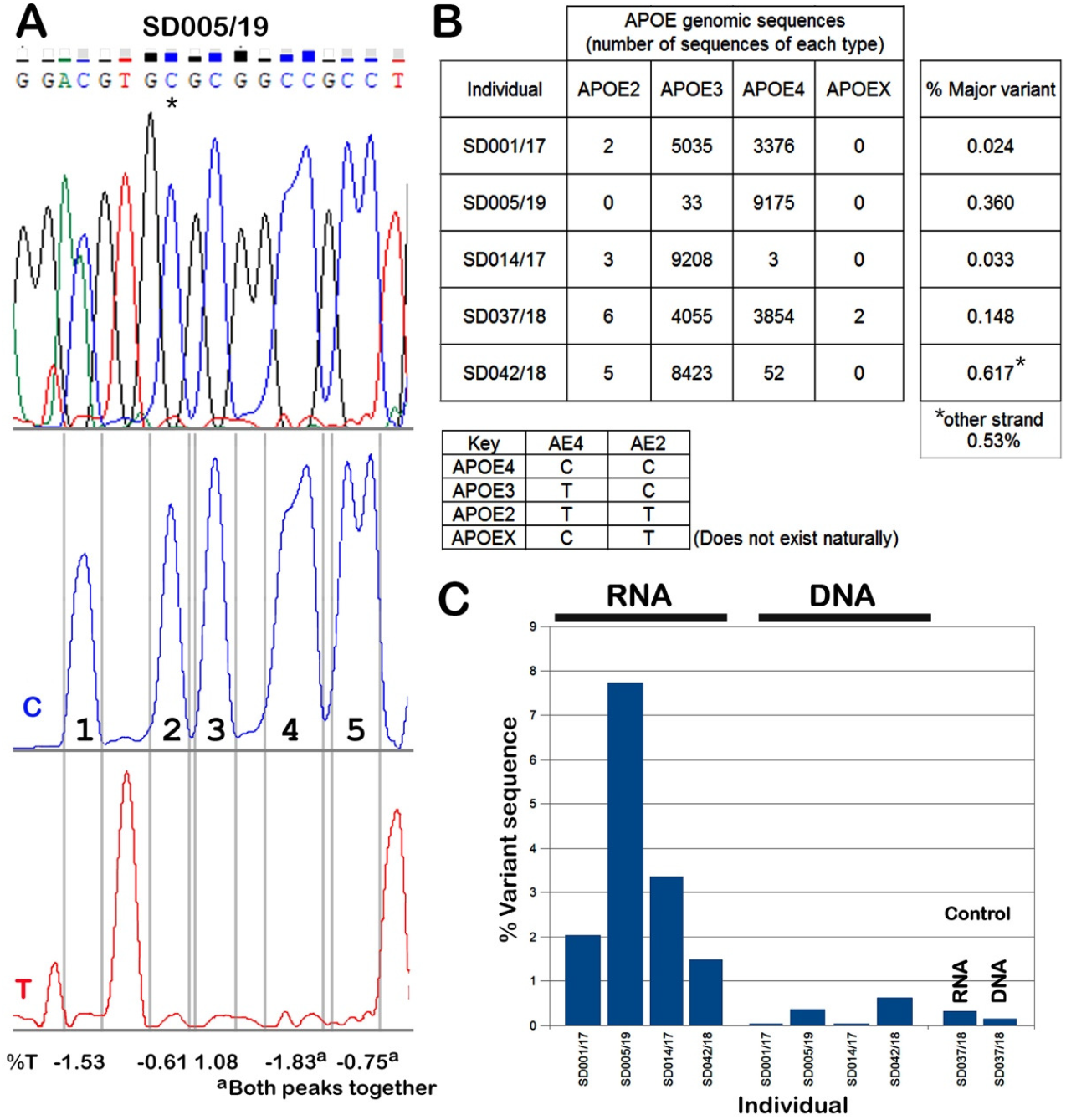
Discordance between transcriptomic and genomic sequences in the key individuals in Fig. 1. (A) Genome: chromatogram of whole-exon sequence data from representative *APOE* genomic DNA (individual SD005/19) showing the location of the AE4 site (asterisk). The % genomic T in at the AE4 position (*) was −0.61% (i.e., zero). (B) Genome: individual molecule sequencing. PCR samples were subjected to single-molecule sequencing, and the numbers of C versus T matches counted and the % variant calculated. (C) Transcriptome versus genome frequencies of each variant in RNA-seq data (left, mean percentages from Table S1) plotted alongside the corresponding frequencies in genomic data (right) for key individuals showing evidence of sequence changes; an individual with no evidence of mRNA sequence changes is shown on the right.

### Brain *APOE* mRNA is abundant in microglia: independent allelic expression in single microglia cells from the same individual

Monoallelic gene expression (MAE) and/or random allelic expression (RAE) are phenomena in which the two alleles of a gene are independently expressed by a single cell. To examine whether MAE/RAE might partly explain our findings, we compared *APOE* mRNA levels in single microglial cells from *APOE* homozygotes and heterozygotes (Fig. 4). mRNA sequence diversification at the AE4 site was exclusively observed in *APOE4*/*E3* heterozygotes, whereas no substantial AE4 diversification was seen in homozygotes. Per-allele relative expression levels in heterozygotes ranged from 0% to 100% (Fig. S1). There was a weak correlation between apparent allele-specific expression and read density, but samples with the highest read densities also showed 0–100% allele-specific expression, which would confirm that *APOE* is subject to MAE/RAE. We also observed up to 8.57% sequence replacements at the AE2 site (Fig. 4), irrespective of zygosity, which cannot be attributed to MAE/RAE because both *APOE4* and *E3* harbor the same nucleotide at this position. Analysis detailed in Fig. 2B further revealed that these changes occur selectively at the AE2 site (Fig. S1). We conclude that MAE/RAE is unlikely to account for the sequence diversification observed in human brain because, in cases where sequence replacements were seen, four of the five individuals are *APOE* homozygotes. In addition, MAE/RAE would not explain the replacements seen in transgenic mice or in cell culture (detailed below) because only one allele is present.

**Fig. 4.**
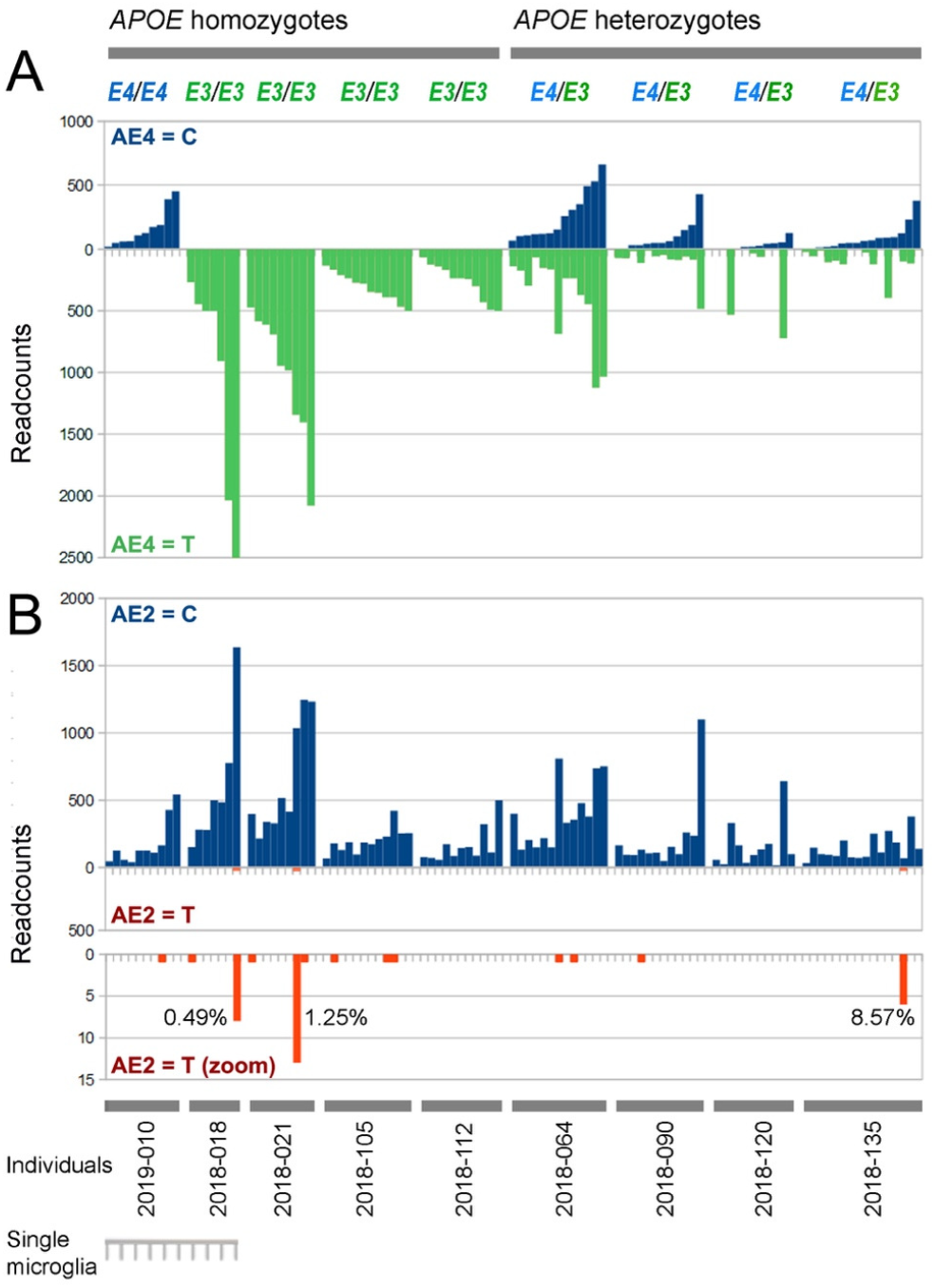
Distribution of *APOE* variants at the AE4 and AE2 sites in single-cell microglia transcriptomic data from nine individuals (primary sequence data reported in (30); the SRAs are given in Table S3). (A) At the AE4 site there were no significant sequence replacements in *APOE* homozygotes, but extensive variation in heterozygotes indicative of monoallelic expression (MAE) or random allelic expression (RAE). (B) At the AE2 site, where all individuals harbor C at this position (either *APOE4* or *E3*) there was evidence of C→T sequence replacements (0.49–8.57%, *P* = <0.01) that were specifically at this site (Fig. S1).

### Site-specific nucleotide replacements in mice expressing human *APOE*

We next addressed whether these sequence changes in *APOE* are seen in a different tissue context. When *APOE4* cDNA was amplified from liver and whole brain of human *APOE4* knock-in mice (17), no C→T changes were found at the AE4 or AE2 sites in mouse liver, but 1 of 48 clones from mouse brain exhibited a C→T change at the AE4 site (Fig. S2). From the sequencing error rate this change would only be expected once in ∼50 experiments of this size (i.e., *P* = 0.02). Two independent RNA-seq datasets from other lines of *APOE* knock-in mice were therefore studied: transgenic mouse brain reported by Lance Johnson and colleagues (18), and brain microglia-like cells reported by Sohail Tavazoie and colleagues (19). Half of the samples (16/30) displayed >0.2% C→T or T→C changes at the AE4 or AE2 sites (values above error rate = 0.045% + 2×SD = 0.165% are significant; *P* = <0.05), and two exceeded 1% (Table S4). There were up to 1.6% C→T replacements the AE4 site in *APOE4* mice, and up to 1.59% reverse T→C replacements in *APOE3* mice. Moreover, in *APOE3* mice there were 0.81% C→T changes at the AE2 site. Only changes at the AE4 and AE2 sites were significant (Fig. S3). Although the frequencies were lower than in human brain, they were statistically significant, with selective nucleotide substitution rates up to 30-fold above the mean error rate. These data suggest that the C→T and/or T→C changes observed in human brain samples are also present in knock-in mice expressing human *APOE*, albeit at a lower frequency.

### APOBEC1 expression replicates the site-specific C→U replacement at the *APOE* AE4 site in human liver-derived cells

The above results suggest that *APOE* transcripts might be subject to RNA editing, the paradigm for which is enzymatic deamination of *APOB* mRNA mediated by APOBEC1 (20). RNA editing of *APOE* has not been reported previously, and we asked whether the canonical RNA-editing enzyme APOBEC1 is active on *APOE* mRNA in cells derived from human liver, a tissue which does not express APOBEC1 (21). Human HepG2 and Huh-7 cells (*APOE4* genotype; Fig. S4) were transiently transfected with either human (Hu) or mouse (Mo) APOBEC1. Both Hu and MoAPOBEC1 were robustly expressed following transfection (Fig. 5A). Amplification and Sanger sequencing of endogenous *APOE* from both genomic DNA and RNA-derived cDNA revealed that both cell lines, transfected with either Hu or MoAPOBEC1, demonstrated >90% C→U RNA editing at the AE4 site and a shift from the *APOE4* genotype to an *E3* ribotype (Fig. 5B,C).

**Fig. 5.**
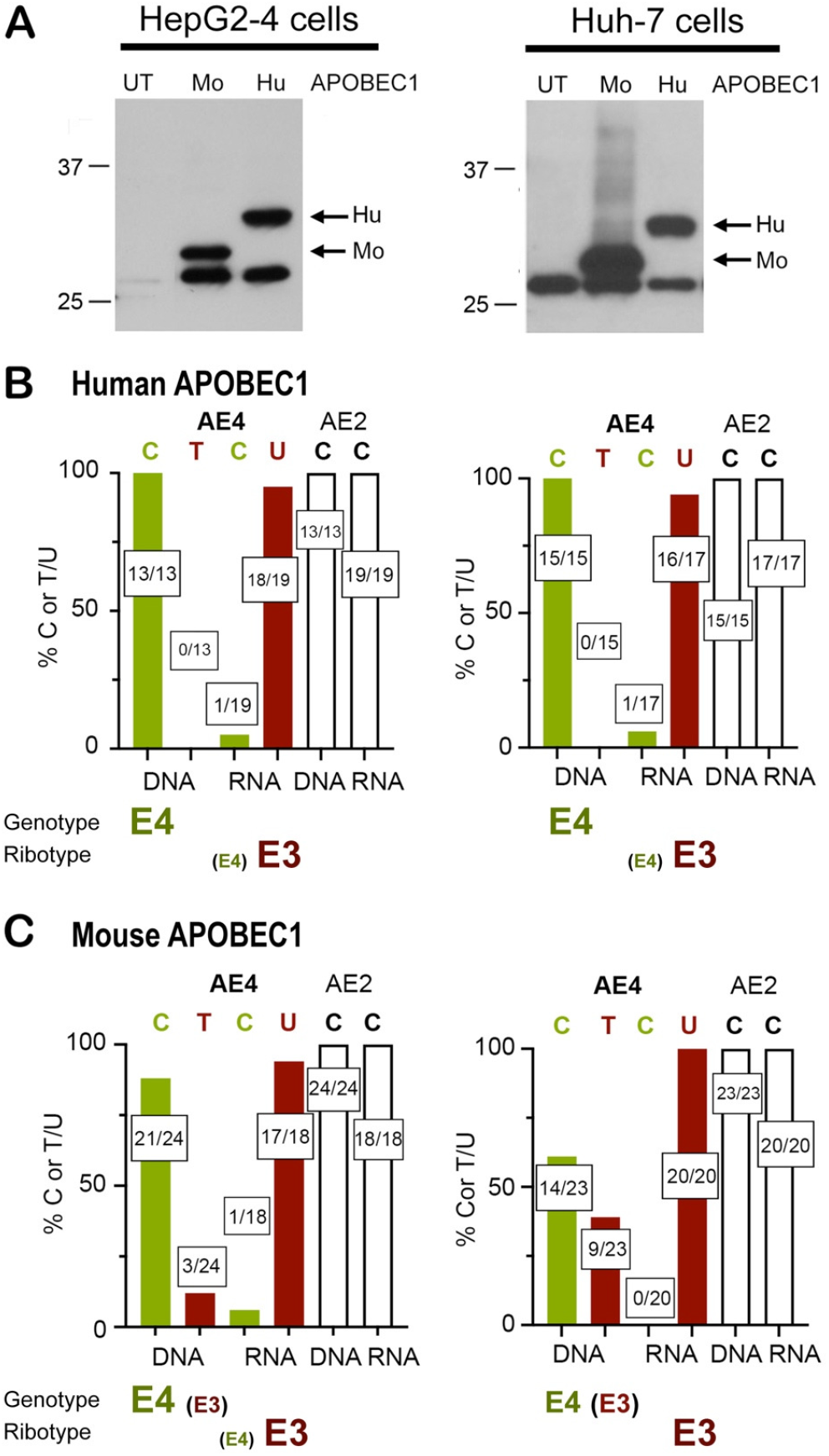
APOBEC1-mediated editing of human *APOE* in cell culture. Human *APOE4* homozygous liver-derived HepG2-4 and Huh-7 cells were transfected with an expression construct for FLAG-tagged human (Hu) or mouse (Mo) APOBEC1. (A) Confirmation of protein expression: transfected cells were analyzed by Western blotting with anti-FLAG antibody; UT, untransfected cells. The band below APOBEC1 protein is nonspecific. (B,C) RNA and DNA from HepG2-4 and Huh-7 cells transfected with either (B) HuAPOBEC1 or (C) MoAPOBEC1 were extracted, cloned, and individual clones were sequenced (Methods). Each graph shows the percentage of C (green) or T/U (red) at the AE4 site, showing that both Mo and HuAPOBEC1 catalyze efficient (>90%) C→U RNA editing at the AE4 site. In addition, MoAPOBEC1 showed evidence of DNA C→T editing at the AE4 site. No changes were observed at any other positions, and there was no evidence of APOBEC-mediated editing of the AE2 site (white) at either the RNA or DNA levels. Boxed numbers indicate the number of clones with C or U/T at a given site over the total number of clones sequenced.

In addition, both lines transfected with MoAPOBEC1 exhibited 12–39% C→T change at the identical AE4 site in genomic DNA (Fig. 5C), whereas no genomic changes were seen with HuAPOBEC1. These findings demonstrate that forced overexpression of APOBEC1 can mediate C→U activity on endogenous *APOE* mRNA at the polymorphic AE4 site, and in the case of MoAPOBEC1 (but not HuAPOBEC1) also mediates C→T changes at the identical site in DNA. These findings add a new dimension to the repertoire of APOBEC1 activity, specifically the demonstration that another endogenous mammalian target (*APOE*) is subject to site-specific C→U RNA editing, and furthermore (with MoAPOBEC1) also undergoes C→T deamination of genomic DNA. Nevertheless, although members of the APOBEC family are obvious candidates, these data do not identify the specific enzyme(s) that might mediate *APOE* editing at the AE4 site *in vivo*, nor do they suggest a plausible candidate for RNA editing at the AE2 site.

### Proteomics reveals non-genomic APOE allelotype polypeptides in human plasma

If edited *APOE* mRNAs are functional *in vivo*, we reasoned that human tissues should contain APOE proteins that differ from their genomically templated versions. To address this we analyzed APOE allelotype variants in publicly available mass spectrometry (MS) proteomic data for 228 human plasma samples (22), specifically focusing on individuals harboring homozygous (either C/C or T/T) genetic allelotypes at either the AE4 or AE2 sites, and determined the proportion of non-genomically encoded peptide forms in the different samples (Figure 6).

**Figure 6.**
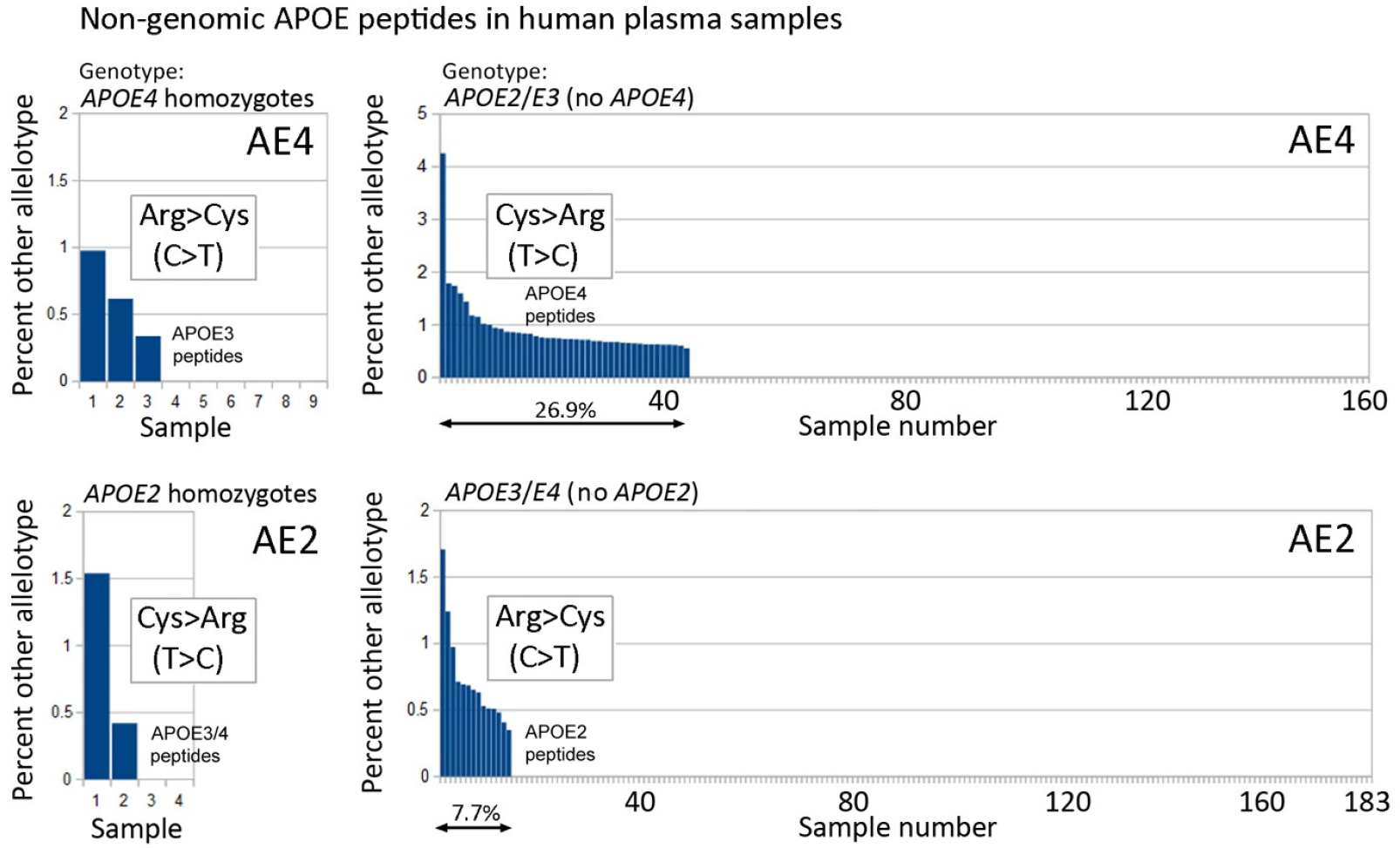
APOE peptides are present in human plasma that are discordant with the underlying *APOE* genotype. The figure plots plasma APOE Arg versus Cys peptide ratios at AE4 (above) and AE2 (below) determined from published datasets of mass spectrometric analysis of samples of different *APOE* genotypes (Methods). Samples were independently sorted left to right in order of abundance of the non-genomic peptide forms.

Multiple individuals were observed to express APOE allelotype peptides that differed from their genotype. Up to 4% of non-genomic APOE4 peptides were present in 43/160 samples (27%) from individuals lacking any *APOE4* allele; these involved Cys>Arg replacements at the AE4 site in APOE polypeptide. Up to 1.7% non-genomic *APOE2* peptides were also present in 14/183 (8%) of individuals lacking any *APOE2* allele; these involved Arg>Cys replacements at the AE2 site. In addition, 3/9 (33%) and 2/4 samples (50%) from *APOE4* and *APOE2* homozygotes, respectively, contained up to 1–1.5% non-genomic peptides that involved Arg>Cys or Cys>Arg replacements at the AE4 or AE2 sites (Figure 1). Although chemical conversion due to peptide or amino acid instability is a potential concern in MS analyses, alternative peptide forms were undetectable in the majority of samples (73% for the AE4 proteotype, 92% for the AE2 proteotype). Because all samples were processed and analyzed using an identical protocol, peptide chemical instability is unlikely to be a contributing factor. Cross-contamination of these samples was also assessed and was found to be negligible (22). The finding of APOE peptides in human plasma that differ from their underlying *APOE* genotype at either the AE4 or AE2 sites therefore substantiates an important prediction of mRNA editing.

There was no evidence of clustering of AE4 and AE2 replacements, and only three samples showed evidence of changes at both positions; all were *APOE3*/*E3* homozygotes with both AE4 Cys>Arg (mRNA T>C, *APOE3*>*APOE4*) and Arg>Cys (C>T, *APOE3*>*APOE2*) replacements. This is not significantly above what would be expected by chance alone (1.33 samples predicted), which again suggests that mRNA editing at the AE4 and AE2 sites is mediated by independent processes.

## Discussion

The central findings of this report are the detection of *APOE3*-like transcripts in the brain of an *E4* homozygous individual, as well as the converse, *APOE4*-like transcripts in individuals who lack an *E4* allele. Our findings emerged from examination of samples of four different regions of human control and AD brain. Genomic DNA sequencing revealed an unexpected divergence in 17 of 39 samples (43.6%) between the genomic and transcriptomic versions of *APOE* in human brain, not only at the AE4 site that discriminates between *APOE4* and *E3*/*E2* but also to a lesser extent at the AE2 site that discriminates between *APOE4*/*E3* and *E2* (Fig. 1).

Several potential confounding factors were considered that might account for the unexpected sequence divergence observed in our human brain transcripts. Cross-contamination and sequencing errors could be excluded (Results), but other possibilities include differences in the transcription or mRNA stability of *APOE4* allelic variants. However, the small difference in per allele mRNA abundance levels, where *APOE4* is expressed at a ∼10–15% higher level than the other alleles (23) could not explain our results. Similarly, monoallelic gene expression (MAE) and/or random allelic expression (RAE) does not provide an explanation because, in cases where sequence replacements were seen in human brain, four of the five individuals are *APOE* homozygotes; in addition, MAE/RAE could not explain the replacements seen in human microglia (AE2), or in transgenic mice (AE4 and AE2) and cell culture (AE4), because only one allele is present.

Low-frequency (10^−6^–10^−7^) single-nucleotide variants are encountered in adult human brain DNA, commonly C→T (24-27), but again would not explain the high frequency of *APOE* variation (up to 10%) observed. Gene duplication could be another potential source of variation, analogous to observations where segments of the *APP* gene undergo somatic gene recombination and reinsertion into the genome via short regions of homology (<20 nt) (28, 29). To address this possibility, transcripts corresponding to full-length *APP* and *APOE* were scanned for single-nucleotide changes. This revealed systematic changes in *APOE*, but none in *APP* (Fig. S5), arguing against this interpretation. In addition, random reintegration of *APOE* transcripts (as reported for *APP*) would preserve the individual allelic type and is unlikely to account for the site-specific variations we report. Other potential confounders could include chimerism/microchimerism and aneuploidy. However, aneuploidies and true chimeras are very rare, and microchimerism (principally through maternofetal exchange) is in the range 0.005% to 0.04% (Extended Discussion 1 in the supplementary material), a rate that is >50-fold lower than the rates of *APOE* sequence divergence observed in the EBB dataset.

Because none of these potential factors provided a satisfactory explanation for the sequence changes in RNA, but not in DNA, other possibilities were considered. First, because *APOE* is abundantly expressed by brain microglia, we interrogated publicly available single-cell RNA-sequencing datasets of independent microglial cells isolated from the brain of the same individual (30). This analysis provided evidence for monoallelic expression (MAE) of APOE, as suggested previously (31, 32). Although no changes were observed at the AE4 site in *APOE* homozygotes, 0.49–8.57% C→U replacements were detected at the AE2 site, findings which cannot be explained by MAE because only one nucleotide is present at this position in the individuals analyzed.

Second, analysis of transcripts from brain and liver of knock-in mice engineered to express human *APOE4* revealed substantial heterogeneity, and 0.42–1.60% C→U changes were seen at the AE4 site in *APOE4* mice, and 0.41–0.81% C→U changes at the AE2 site. In addition, 0.44–1.59% U→C changes were seen at the AE4 site in *APOE3* mice, mirroring similar changes observed in human brain from *APOE3* homozygous individuals, albeit at a lower frequency.

Having ruled out alternative explanations, a remaining interpretation is that the differences between the APOE transcriptomic sequence and the underlying genomic sequence reflect RNA editing. The paradigm for C→U RNA editing is cytosine deamination of *APOB* RNA mediated by the APOB mRNA editing enzyme APOBEC1 in conjunction with the RNA-binding proteins RBM47 and A1CF (33-35). U→C transitions are also common (36), which would indicate that some enzymes can operate bidirectionally (Extended Discussion 2 in the Supplementary Material), although to our knowledge this has not yet been demonstrated for APOBEC family members. To directly address the possibility that *APOE* mRNA is subject to RNA editing, we transfected human or mouse APOBEC1 into cultured human liver-derived cells expressing *APOE4*. This led to efficient (>90%) specific replacement of the C nucleotide at the AE4 site of *APOE4* mRNA by U (T in cDNA), in two different cell lines, in the absence of changes at other sites including AE2. APOBEC1-dependent editing of the same nucleotide in the underlying genomic DNA was also found, but only with mouse (not human) APOBEC1 (12–39%). These findings demonstrate that human *APOE4* mRNA can undergo C→U RNA editing under circumstances when APOBEC1 is overexpressed, and at the specific nucleotide that differentiates the *APOE4* sequence from *E3* and *E2*. Nevertheless, APOBEC-mediated C→U RNA editing of *APOE* in brain microglia remains to be formally demonstrated, and the enzyme(s) and/or cofactors involved in modifying the AE2 site are unknown.

To determine whether edited mRNA transcripts are functional *in vivo*, we examined human plasma proteomic data. This confirmed the presence of APOE peptide allelic variants that differ from the underlying *APOE* genotype (Fig. 6). The observed peptide changes were most frequently observed at the AE4 site and to a lesser extent at the AE2 site, findings that replicate the patterns of mRNA editing found in human brain and in *APOE* transgenic mice. Similar proteotype/genotype discordances have been noted previously. In one study the *APOE* allelotypes of 7/164 individuals were misclassified from plasma proteomic analysis, even though their DNA-based genotypes were carefully reconfirmed (37), and in another study 3/172 individuals were misclassified from plasma data despite DNA sequencing to establish their genotypes (22). However, no hypotheses were advanced to explain these sequence discordances. The proteomic data thus provide independent support for the suggestion that human *APOE* mRNA editing occurs *in vivo*. However, these findings leave unanswered the tissue location and cell-type responsible because *APOE* is expressed widely, with high levels in several tissues as well as in brain, all of which can exchange with the blood. Plasma APOE proteomes may therefore provide a whole-body estimate of the extent of mRNA editing, which may be locally variable among tissues and between different individuals.

Several aspects of our findings merit further discussion. First, RNA editing was only observed in six of nine individuals, and some individuals showed prominent editing whereas others showed none. There were no statistically significant correlations between RNA editing and sex, age, post-mortem interval, disease status, or Braak and Thal staging (not presented). Nevertheless, of the four individuals with the highest levels of RNA editing, three had been diagnosed with AD. In our microglia data the highest level of editing was in an individual diagnosed with AD. Given that APOBEC enzymes are inducible by cytokines/infection/inflammation (38-41), it is possible that editing is induced by local pathological changes in brain; studies in extended cohorts will be essential to confirm and extend our findings.

Second, although APOBEC1 is able to catalyze *APOE* AE4 nucleotide substitution, this does not identify the specific enzyme responsible *in vivo* because APOBEC1 is only one of a large family of APOBEC-related enzymes, several of which can mediate C→U changes in RNA (42-45). At least eight different APOBECs are expressed in our brain samples (Figure S8), but formal identification of functionality is complicated because of highly homologous sequences, alternative splicing, and variable expression. In addition, there were no replacements at the AE2 site with APOBEC1 *in vitro*, and we infer that substitutions at the two sites require different enzyme(s) and/or cofactor stoichiometries within the editosome (34). Other potential enzymatic modification pathways that might explain our findings were also considered, including editing by activation-induced deaminase (AID), replacements driven by CpG methylation or RNA methylation, and ADAR (adenosine deaminase acting on RNA)-mediated A→I (i.e., A→G) changes in antisense RNA. However, these possibilities could be ruled out (Extended Discussion 3 in the supplementary material, also Figs S5 and S6).

A further outcome relates to divergence in the genotype/ribotype of *APOE* in human liver-derived cells. Based on earlier reports that two commonly used liver cell lines (HepG2, Huh-7) are *APOE3*/*E3* (46, 47), we analyzed *APOE* genotype and ribotype from four different clones of HepG2 cells from the same source (ATCC, Methods) which have been maintained over more than 20 years, along with several other human liver-derived cell lines. Among the four different HepG2 clonal lines, and contrary to our expectations, all exhibited predominantly (clones 1–3) or exclusively (clone 4) an *APOE4*/*E4* genotype (Fig. S4). These findings suggest that investigators using human liver-derived cell lines for studies of APOE function should consider validating the *APOE* genotype/ribotype.

This is the first report of *APOE* mRNA editing, and our conclusions must remain tentative until they can be validated in other cohorts by independent investigators. Neverthess, we report site-specific changes not only in human brain RNA but also in human microglia and in *APOE* transgenic mice. Moreover, we report efficient APOBEC1-mediated *APOE* mRNA editing in *vitro*. Finally, we provide evidence that peptides corresponding to edited APOE mRNAs are present in human plasma. If confirmed, these data potentially afford a new perspective on the biology of human *APOE*.

## Acknowledgments

We would like to express our sincerest gratitude to all researchers who filed their data online that permitted investigation of the wider generality of the phenomenon described in this paper. We thank Colin Smith and Chris-Anne Mckenzie at the Edinburgh Brain Bank for providing samples for deep sequencing. We thank Nanda Kumar and Rudy Tanzi (MassGeneral Institute for Neurodegenerative Disease and Harvard Medical School) for discussions and confirming sequence changes in some of their samples. James Lathe is thanked for help with statistical analyses. The primary brain sequencing study was funded by the Benter Foundation (RL and JH); NOD and VB were supported by grants from the National Institutes of Health (DK-52574, DK-119437, and HL-151328).

## Author contributions

Conceptualization: RL, JH, NOD. Methodology: VB, SJG, JH, NOD, RL. Investigation: VB, SJG, JH, NOD, RL. Funding acquisition: RL, JH, NOD. Project administration: RL, JH, NOD. Writing – original draft: RL, NOD, VB, SJG. Writing – review & editing: VB, SJG, JH, NOD, RL.

## Declaration of interests

The authors declare no competing interests.

## Supplementary materials

Supplementary data associated with this article can be found, in the online version, at DOI XXX {DOI to be inserted}.

## Methods

### RNA sequencing and datasets

Deep sequencing of RNA from human brain was described previously (15, 16). These datasets have been uploaded to the National Institute of Biotechnology Information (NCBI) and are freely available online (SRA numbers are listed in Table S1). Microglial RNA-seq datasets from the Molecular Neurobiology Section at the University of Groningen, The Netherlands (Bioproject PRJNA611563) (30) filed at NCBI were also analyzed (details in Tables S1 and S2). The genotypes of the donor individuals were kindly provided by Bart Eggen (Groningen) and Isabell Ehmer at the Netherlands Brain Bank under project agreement NBB 1855. For mice, RNA-seq data of brain from human *APOE* transgenic mice filed online at NCBI by the University of Kentucky (Bioproject PRJNA875087) (18) and by Rockefeller University, USA (Bioproject PRJNA1001862) (19) were analyzed (details in Table S4).

### Analysis of allelic variants

Analyses involved BLAST searching at NCBI with either a cutoff of a maximum of 100 matching sequences (for long probes) or a cutoff of 5000 matches (for short probes). The two types of probes employed give slightly different results. Long probes (64-mers) detect both perfect matches and mismatches and require manual filtering, whereas 36-mer probes only detect perfect matches using BLAST default settings. Long-probe BLAST displays the perfect matches first (e.g., Fig. 1B), followed by allelic variants (located centrally), short transcripts, and sequencing errors (non-central). Probe sequences and further details are given in the Supplementary Methods.

### PCR amplification of the genomic *APOE* locus from brain and DNA sequencing

Brain DNA was extracted fusing the QIAGEN DNeasy Blood and Tissue Mini kit and the central region of *APOE* exon 4 was PCR amplified using the high-fidelity KOD polymerase kit (SigmaAldrich) and subjected to direct population Sanger sequencing (GATC Biotech/Eurofins); areas under the chromatograms were then quantified using ImageJ. For individual molecule sequencing, DNA library preparations were performed by Genewiz (Leipzig, Germany) using the NEBNext Ultra II DNA Library Prep kit and sequenced (Illumina) (details in Supplementary Methods).

### Animals and cell lines

Human *APOE4* knock-in mice (16 weeks old) were kindly provided by Dr David Holtzman, Department of Neurology, Washington University School of Medicine, St Louis. Different lines of human HepG2 cells obtained from the ATCC (HB-8065); lines HepG2-1 and 3 originated from 2025 (respective numbers of passages 24 and 20), HepG2-2 (1996, number of passages 56), and HepG2-4 from 1995 (number of passages 53). Human Huh-7 cells were from the JCRB Cell Bank (JCRB0403) and the mouse hepatocyte cell line AML12 was from ATCC (CRL-2254) (details in Supplementary Methods).

### Ribotyping of transgenic mice tissues

Total RNA from liver and brain from five human *APOE4* knock-in mice (2 males and 3 females) was DNAse treated, the region encompassing the AE4 and AE2 sites was PCR amplified, cloned, and individual clones were submitted to Azenta Life Sciences/Genewiz for Sanger sequencing (details in Supplementary Methods).

### *In vitro* editing: APOBEC transfection, ribotyping, and genotyping

Cells were transiently transfected with a eukaryotic pCMV vector expressing either FLAG-tagged murine or human APOBEC1. Forty-eight hours after transfection, DNase-treated RNAs were used to synthesize cDNA and the *APOE* region containing the AE4 and AE2 sites was PCR amplified using Platinum SuperFi DNA polymerase (Invitrogen), gel purifed, cloned, and individual clones submitted for Sanger sequencing as before. Genomic DNA was prepared by RNase A and proteinase K treatment and isopropanol precipitation. Cloning and sequencing of individual clones was as before (details in Supplementary Methods).

### Protein extraction and western blotting

Washed cells were lyzed, cleared by centrifugation, resolved by SDS-PAGE, transferred to polyvinylidene fluoride (PVDF) membranes, and probed with rabbit anti-FLAG antibody (Invitrogen).

### Proteomic analysis of human plasma

Mass spectrometric data for the abundance of APOE peptides in 228 samples (including 12 replicates) of plasma obtained from individuals participating in the Gait and Brain Study in Canada (48) were retrieved from the supplementary material filed online at https://pubs.acs.org/doi/10.1021/acs.jproteome.3c00557 (22). Gender, age, and disease status data for these specific individuals were not available. APOE proteotype analysis is facilitated because both the amino acid replacements that distinguish between APOE4, E3, and E2 proteins (Cys↔Arg at AE4 and AE2) alter tryptic cleavage of the polypeptide. For the AE4 site the abundance was studied of tryptic peptide analyte B ({R}LGADMEDVR), that is APOE4-specific, and analyte A ({R}LGADMEDVCGR), that is present in both APOE3 and E4 proteins. For the AE2 site, the abundance was studied of analyte C ({R}LAVYQAGAR), that is present in both APOE3 and E4, and analyte D ({K}CLAVYQAGAR), that is APOE2-specific. The A/B and C/D peptide ratios were calculated based on the values for peptide intensities normalized to the internal standard analyte ({R}LGPLVEQGR; ’area ratio’ in column E of the online supplementary table in file pr3c00557_si_002.xlsx) (22). To evaluate potential interconversion of peptide allelotypes at the AE4 at AE2 sites, A/B and C/D ratios were determined separately in samples from individuals who were homozygous at either the AE4 or the AE3 sites. A ratio of, for example, 1 to 100 for the genomically absent proteotype versus the dominant proteotype was taken to indicate ∼1% *in vivo* conversion.

## Notes

### Competing Interest Statement

The authors have declared no competing interest.

