## Supplementary data for "mRNA editing of the Alzheimer’s risk gene *APOE*"

- **Extended Methods**
- **Extended Discussion 1.** Chimerism/microchimerism and aneuploidy.
- **Extended Discussion 2.** Wider evidence for both C→U and U→C editing.
- **Extended Discussion 3.** Activation-induced deaminase (AID), CpG methylation, RNA methylation, ADAR (adenosine deaminase acting on RNA), and sequence-specific mutations.
- **Table S1.** Underlying genotypes, individuals, brain samples, datasets, and numbers of matches to AE4 and AE2 C versus T probes: primary data.
- **Table S2.** Analysis of single-cell RNA sequencing data from brain microglia from control and Alzheimer's disease (AD) individuals.
- **Table S3.** List of single-cell microglia datasets analyzed in Figure 4 in the main text.
- **Table S4.** Primary *APOE* sequence variation datasets for transgenic mice expressing either human *APOE4* or *APOE3*.
- **Figure S1.** Specificity of sequence changes at the AE2 site in individual microglia, and variation in per-cell expression levels.
- **Figure S2.** Studies on *APOE* ribotype in transgenic mice expressing human *APOE4*.
- **Figure S3.** Non-genomic transcripts in transgenic mice expressing human *APOE4*.
- **Figure S4.** Unexplained variation in *APOE* genotype and ribotype in human liver-derived cells.
- **Figure S5.** Screening of the entire *APOE* and *APP* transcripts for single-nucleotide changes.
- **Figure S6.** Distribution of CpG nucleotides in *APOE* cDNA.
- **Figure S7.** Analysis of potential antisense transcription of human *APOE*.
- **Figure S8.** *APOBEC* mRNA expression in human brain.

### Extended Methods

#### RNA sequencing and datasets

Deep sequencing of RNA from human brain (RNA-seq) was described previously (Hu *et al.* 2022; Hu *et al.* 2023). These datasets have been uploaded to the National Institute of Biotechnology Information (NCBI) and are freely available online (SRA numbers are listed in Table 1). NCBI microglial RNA-seq datasets from the Molecular Neurobiology Section at the University of Groningen, The Netherlands (Bioproject PRJNA611563) (Alsema *et al.* 2020) were also analyzed (details in Tables S1 and S2). The genotypes of the donor individuals were kindly provided by Bart Eggen (Groningen) and Isabell Ehmer at the Netherlands Brain Bank under project agreement NBB 1855. For mice, RNA-seq data of brain from human *APOE* transgenic mice filed online at NCBI by the University of Kentucky (Bioproject PRJNA875087) (Lee *et al.* 2023) and by Rockefeller University, USA (Bioproject PRJNA1001862) (Millet *et al.* 2024) were analyzed (details in Table S4).

#### Analysis of allelic variants

Analyses involved BLAST (Altschul *et al.* 1990) searching at NCBI with either a cutoff of a maximum of 100 matching sequences (for long probes) or a cutoff of 5000 matches (for short probes). The two types of probes employed give slightly different results. Long probes (64-mers) detect both perfect matches and mismatches and require manual filtering, whereas 36-mer probes only detect perfect matches using BLAST default settings. BLAST displays the perfect matches first (e.g., Figure 1B), followed by allelic variants (located centrally), short transcripts, and sequencing errors (non-central). Of note, automated RNA-editing analysis programs and associated databases were not used for two reasons. First, they principally (but not exclusively) focus on A–I rather than C–U editing. A more severe limitation is that, following the initial BLAST-like algorithm to compare mRNA and genomic sequences, they remove known SNPs from the list of potential editing sites. Both constrain analysis of the two *APOE* sites.

#### APOE probes (long versions; 64-mers)

>AE4_C=APOE4
GGCCCGGCTGGGCGCGGACATGGAGGACGTG**C**GCGGCCGCCTGGTGCAGTACCGCGG
CGAGGTG
>AE4_T=APOE2+APOE3
GGCCCGGCTGGGCGCGGACATGGAGGACGTG**T**GCGGCCGCCTGGTGCAGTACCGCGG
CGAGGTG
>AE2_C=APOE3+APOE4
GCTCCTCCGCGATGCCGATGACCTGCAGAAG**C**GCCTGGCAGTGTACCAGGCCGGGGCC
CGCGAG
>AE2_T=APOE2
GCTCCTCCGCGATGCCGATGACCTGCAGAAG**T**GCCTGGCAGTGTACCAGGCCGGGGCC
CGCGAG

#### APOE probes (allele-specific; 36-mers)

>AE4_E4=C
CGGACATGGAGGACGTG**C**GCGGCCGCCTGGTGCAGT
>AE4_E23=T
CGGACATGGAGGACGTG**T**GCGGCCGCCTGGTGCAGT
>AE2_E34=C
CCGATGACCTGCAGAAG**C**GCCTGGCAGTGTACCAGG
>AE2_E2=T
CCGATGACCTGCAGAAG**T**GCCTGGCAGTGTACCAGG

**PCR amplification of the genomic *APOE* locus from brain and DNA sequencing**
DNA was extracted from brain (cerebellum) samples from key Edinburgh Brain Bank individuals using the QIAGEN DNeasy Blood and Tissue Mini kit involving protease K treatment at 55°C. The central region of APOE exon 4 (260 nt) was denatured (95°C, 2 minutes) and PCR amplified using primers CAC GGC TGT CCA AGG AGC TGC (upstream) and GAT GGC GCT GAG GCC GCG CTC (downstream) using the high-fidelity KOD polymerase kit (SigmaAldrich) and a gradient PCR cycle {elongation (54.3–66.6°C for 20 s) followed by 10 s at 70°C, and denaturation (20 s at 95°C)}. PCR products were run on a 1.5% agarose gel, and the DNA bands were recovered and purified using the QIAGEN extraction kit. The PCR products were subjected to direct population Sanger sequencing (GATC Biotech/Eurofins), the sequence data generated were visualized in CHROMAS (Technelysium DNA Sequencing Software), and the areas under the chromatograms were then quantified using ImageJ (Schneider *et al.* 2012).

For individual molecule sequencing, DNA library preparations were performed by Genewiz (Leipzig, Germany) using the NEBNext Ultra II DNA Library Prep kit following the manufacturer's recommendations. Briefly, end-repaired adapters were ligated after adenylation of the 3′-ends followed by enrichment by limited cycle PCR. DNA libraries were validated and quantified before loading. The pooled DNA libraries were loaded on the Illumina instrument according to manufacturer's instructions. The samples were sequenced using a 2 × 250 paired-end (PE) configuration. Image analysis and base calling were conducted using Illumina Control Software on the Illumina instrument. The raw Illumina reads were checked for adapters and quality via FastQC. The raw Illumina sequence reads were trimmed of their adapters using Trimmomatic v. 0.36. Raw sequence data (.bcl files) generated from Illumina MiSeq were converted into fastq files and demultiplexed using the Illumina bsl2fastq v. 2.17 program.

**Cell lines and animals**
Human *APOE4* knock-in mice (16 weeks old) were kindly provided by Dr David Holtzman, Department of Neurology, Washington University School of Medicine, St Louis. Different lines of human HepG2 cells obtained from the ATCC (HB-8065) were maintained in Gibco Minimum Essential Medium supplemented with 10% fetal bovine serum (FBS) and penicillin/streptomycin. Lines HepG2-1 and 3 originated from 2025 (respective numbers of passages 24 and 20), HepG2-2 from 1996 (56 passages), and HepG2-4 from 1995 (53 passages). Human Huh-7 cells (JCRB Cell Bank JCRB0403) were grown in Gibco Dulbecco's modified Eagle medium (DMEM/F12) in the presence of 10% FBS and penicillin/streptomycin (number of passages = 21). The mouse hepatocyte cell line AML12 (ATCC: CRL-2254) was maintained in DMEM/F12 medium supplemented with 10% FBS, 10 mg/ml insulin, 5.5 mg/ml transferrin, 5 ng/ml selenium, 40 ng/ml dexamethasone, and penicillin/streptomycin.

**Ribotyping of transgenic mice tissues**
Total RNA from liver and brain from five human *APOE4* knock-in mice (2 males and 3 females) was prepared using TRIzol (Invitrogen) following the manufacturer's protocol. Isolated total RNA (10 µg) was treated with Turbo DNAse from the Turbo DNA-free kit and DNA synthesis was performed using random primers and the MultiScribe high-capacity cDNA reverse transcription kit (Applied Biosystems). cDNA was used to amplify a 827 bp region of human *APOE4* encompassing the AE4 and AE2 sites. PCR reactions were performed with the following human-specific *APOE4* primers: Fwd HuE4-85 GGA GCC CGA GCT GCG CCA G and Rev HuE4-912 GGC AGC CTG CAC CTT CTC CAC CAG CCC G. Cycling conditions were 98°C 2 minutes, 37 cycles at 98°C/30 s, 72°C/30 s, 72°C/50 s, followed by 72°C for 10 minutes. The PCR product was gel purifed on a 1.4% agarose gel and cloned into the pCR-Blunt II-TOPO vector (Invitrogen) following the manufacturer's recommendations. Individual clones were isolated and submitted to Azenta Life Sciences/Genewiz for Sanger sequencing.

***In vitro* editing: APOBEC transfection, ribotyping, and genotyping**
Cells were grown to 50–60% confluence and transiently transfected with 3 µg of a eukaryotic pCMV vector expressing either FLAG-tagged murine or human APOBEC1 using Lipofectamine 3000 (Invitrogen). Forty-eight hours after transfection, DNase-treated RNAs were used to synthesize cDNA as described above, and the *APOE* region containing the AE4 and AE2 sites was amplified using Platinum SuperFi DNA polymerase (Invitrogen) with the following primers: Fwd GGT CAC CCA GGA ACT GAG G and Rev GGG CTC GAA CCA GCT CTT GAG GC, resulting in a 636 bp PCR product. Cycling conditions were 98°C 2 minutes, 37 cycles at 98°C/30 s, 65°C/30 s, 72°C/50 s, and 72°C for 10 minutes. The PCR product was gel purified, cloned, and individual clones were isolated and submitted for Sanger sequencing as before. Genomic DNA was prepared as follows: following two washes in PBS, cells were scraped into cell lysis buffer supplemented with 2 µl RNase A (10 mg/ml; QIAGEN) and incubated at 37°C for 30–60 minutes. Proteinase K (3 µl; 20 mg/ml) was added and the cell lysate was incubated at 55°C for 2–3 h. Proteins were precipitated from the lysate and DNA was recovered by isopropanol precipitation. After a 70% ethanol wash, DNA was air-dried for 15 minutes, resuspended in water, and rehydrated overnight before cloning and sequencing of individual clones as described above.

**Protein extraction and western blotting**
Cells were washed with PBS and resuspended in 70 ml of RIPA buffer containing 25 mM Tris (pH 7.4), 0.15 M NaCl, 1% NP40, 1% sodium deoxycholate, 1 mM EDTA, 1 mM sodium vanadate, 0.1 % SDS, 50 mM β-glycerophosphate, 100 mM sodium fluoride, and a 1× concentration of protease inhibitor (Roche Applied Science). Lysis was completed by three passages through an 18-gauge needle. After incubation for 15 minutes on ice, the lysate was cleared by centrifugation at 15,000 g for 10 minutes at 4°C. Aliquots of homogenate (20 µg) were resolved by 10% SDS-PAGE, transferred to polyvinylidene fluoride (PVDF) membranes, and probed with rabbit anti-FLAG antibody (Invitrogen) at 1:1000 dilution.

### Extended Discussion 1. Chimerism/microchimerism and aneuploidy

We carefully considered whether aneuploidy and/or chimerism might potentially account for our findings. Although trisomy of chromosome 19 (bearing *APOE*) is reported in patients with chronic myeloid leukemia (Rousseau *et al.* 2007), none of our individuals suffered from this condition. Mosaic trisomies arising through chromosome missegregation during development are also extremely rare (<1/10 000 live births). For completeness, the possibility was considered that some individuals might be chimeras of different *APOE* genotypes. However, full chimeras in human are rare (1 in 1000 live births) (Madan 2020) and are unlikely to explain the common AE4 diversification we observe.

Microchimerism through maternofetal exchange is more plausible because maternal and fetal pluripotent stem cells can contribute to multiple tissues (Klonisch & Drouin 2009). Persistence of maternal cells is reported in mouse brain microglia, endothelial, and neuron-like cells (Schepanski *et al.* 2022), and examination of human adult male brain tissues for female-specific (maternal) markers revealed 0–9 female cells per 4000–8000 male cells (mean 0.04%) (Snethen *et al.* 2020). However, this rate is >50-fold lower than the rates of *APOE* variant divergence observed in the EBB dataset. For persistence of fetal cells, Chan *et al.* reported that 64/183 (35%) of human female brain specimens examined tested positive for male-specific genetic markers, likely emanating from residual fetal male cells acquired while gestating a male fetus (Chan *et al.* 2012). However, the extent of chimerism was extremely low, with a maximum contribution of 0.005%, far below the extent of replacement we observe. In addition, even allowing for female/female in addition to female/male chimerism, the overall frequency is far less than the frequency of individuals showing an abnormal ratio at the AE4 site (up to one third of individuals). Based on these considerations, microchimerism is unlikely to explain the results.

---

### Extended Discussion 2. Other evidence for both C→U and U→C editing.

APOBEC enzymes are generally thought to catalyze deamination, but we found both C→U and U→C transitions, which would imply that some of these enzymes might catalyze the reverse (reamination) reaction. Indeed, C→U deamination is inherently reversible (Cohen & Wolfenden 1971). Although C→U conversion is the canonical reaction, U→C conversions have been widely reported. Selective U→C replacments are reported in *WT1* (Sharma *et al.* 1994) and *TPH2* (Grohmann *et al.* 2010) mRNAs, and further U→C changes have been confirmed elsewhere in *WT1* mRNA where antisense A→I editing could be excluded (Niavarani *et al.* 2015). U→C editing has also been reported in *ADAR2* transcripts (Laxminarayana *et al.* 2007). In bovine mammary gland, U→C changes are more common than C→U (Lopdell *et al.* 2019), and U→C changes are also as common as C→U replacements in mouse spermatogenesis (Wang *et al.* 2019). In addition, the human BigBrain project (Dredge *et al.* 2025) reported several instances of U→C editing that could not be attributed to A→G editing in the other strand.

Although any one of these reports might be open to challenge, the combined weight of evidence argues that U→C editing does take place in mammalian cells. This has been confirmed by proteomic analysis of human B cells, where several peptides predicted from U→C RNA editing of other editing events were detected by mass spectrometry (Li *et al.* 2011), as for APOE, which would indicate that these are true editing events taking place *in vivo*, although the editing machinery involved was not identified.

Which specific enzymes might be responsible? We have demonstrated C→U RNA editing at the AE4 site of *APOE* by APOBEC1, but this enzyme is poorly expressed in brain, although it cannot be formally excluded as a candidate (Figure S8). However, APOBEC3A, APOBEC3G, and APOBEC3B also have C→U editing activity (Sharma *et al.* 2015; Sharma & Baysal 2017; Alonso de la Vega *et al.* 2023; Zhang *et al.* 2024), and, given their close evolutionary relationships, other

APOBECs may also be capable of catalyzing this conversion, although this remains to be demonstrated. Furthermore, there is evidence that some sites are targets for more than one APOBEC enzyme. For example, Kim *et al.* report multiple sites in the genome of the RNA virus SARS-CoV-2 that are independently edited by both APOBEC1 and APOBEC3A, and some that are edited by both APOBEC3A and APOBEC3G (Kim *et al.* 2022). Other important considerations are (i) homo- and heterodimerization modulate APOBEC editing activity (heterodimerization of APOBEC3G with APOBEC3F stimulates its editing activity (Ara *et al.* 2017)), and (ii) some APOBEC enzymes are dependent on cofactors such as A1CF. Our preliminary data (not presented) indicate that APOBEC3G editing is prevalent in brain and pairwise correlation analysis reveals that *A1CF* expression is associated with *APOE* editing ($P$ = 0.03). However, these are preliminary findings that warrant detailed analysis.

It is very likely that some APOBECs may also catalyze the reverse U→C transition because, in HIV-1-infected cells treated with interferon-α to specifically induce APOBEC activity, U/T→C transitions were as common as C→U transitions (Koning *et al.* 2011), although both DNA and RNA may have been targeted. It is therefore likely that some APOBEC enzymes can operate bidirectionally, although this remains to be confirmed through studies using purified enzymes, a topic for future research.

Of interest, APOBEC3A (which is expressed in brain; Figure S8) is apparently capable of catalyzing G→A RNA editing in addition to canonical C→U editing (Niavarani *et al.* 2015), which the authors attributed to re-amination of G to 2,6-diaminopurine (an analog of A); this argues that some enzymes can be promiscuous in the specific reactions they catalyze. This is supported by the diversity of substitutions attributed to editing: in addition to canonical APOBEC-mediated C→U/T and ADAR-mediated A→G conversions, other replacements including U/T→C, C→A, G→A and others have been validated (Pickrell *et al.* 2012; Peng *et al.* 2012; Guo *et al.* 2019), but the specific enzymes (or enzyme and cofactor combinations) responsible have not yet been identified.

---

### Extended Discussion 3. Activation-induced deaminase (AID), CpG methylation, RNA methylation, ADAR (adenosine deaminase acting on RNA), and sequence-specific mutations

Other potential enzymatic modification pathways that might explain our findings were also considered. Immunoglobulin variable regions undergo somatic hypermutation to expand antibody diversity. Mutations are triggered by activation-induced deaminase (AID), an APOBEC-related enzyme (Muramatsu *et al.* 1999) that preferentially deaminates cytosine in DNA residues, leading to C→T transitions (Di Noia & Neuberger 2007), although error-prone repair mechanisms can generate many other replacements (reviewed in (Pilzecker & Jacobs 2019)). However, in contrast to the wide spread of AID-dependent sites of mutation across a whole region of the target DNA, we observed highly site-specific changes in *APOE*.

In addition, methylation of cytosines in DNA makes them prone to spontaneous and enzymatic deamination that generates thymine residues. Both inferred editing sites are within a CpG nucleotide-rich locus, where the ancestral *APOE4* is C and the human-specific versions E3 and E2 represent stepwise replacement of CpG by TpG. Although *APOE* is CpG-methylated in brain (Yu *et al.* 2013), and there are potential mechanisms to convert 5-methylcytosine (5mC) to T, only the AE4 and AE2 sites are altered, even though the entire *APOE* locus is rich in CpG methylation sites (Figure S6), suggesting that DNA methylation itself would not explain the specific C→T transitions (nor the observed T→C changes).

RNA methylation is a further potential mechanism. Although several studies have mapped 5mC sites in human mRNAs and non-coding RNAs, no sites were found in *APOE* transcripts (Squires *et al.* 2012; Huang *et al.* 2019).

We also considered the possibility of RNA editing mediated by ADAR (adenosine deaminase acting on RNA). Although this enzyme catalyzes A→I (i.e., A→G) changes, in contrast to the prominent C→U/T changes we observe, ADAR-mediated editing of antisense RNAs can result in apparent U/T→C base changes in overlapping sense transcripts/cDNAs (Pecori *et al.* 2022). Based on a single earlier report that an antisense transcript (AS1) exists for human *APOE* (Seitz *et al.* 2005), we screened our RNA-seq libraries for potential antisense transcripts: this set an upper limit of 1.34% for antisense transcription (Figure S7). This low level would not explain the extent of sequence changes (up to 10% in brain, and up to 100% *in vitro*).

In addition, because sequencing misreads tend to occur at the ends of sequencing reads (Pickrell *et al.* 2012), we examined the observed mismatches for their location with each read. As shown in Figure 1 in the main text and Figure S4 below, the mismatches are centrally placed within the reads, which would exclude end-associated misreads as an explanation.

Finally, sequence-specific mutations either *in vivo* or introduced during the reverse transcription and sequencing steps could potentially introduce a further complexity. Both sites center on the trinucleotide GCG, and we therefore addressed whether this sequence might be specifically prone to alteration. Several observations militate against. In addition to the AE4/AE2 sites, full-length *APOE* and *APP* (Figure S5, below) mRNAs contain 45 and 29 GCG sites, respectively, but with no evidence of selective nucleotide replacements. We also examined 50 kb of well-expressed brain transcripts specifically for substitution rates in 3-mer, 4-mer, 5-mer, and 6-mer nucleotide sequences identical in context to the GCG motifs in *APOE*. The frequency of changes was no different from the overall rate of replacements listed in Figure 2C. Furthermore, sequence-specific changes during work-up are also inconsistent with the finding almost 10% changes in some samples, but 0% in others, using identical technology. Selective sequence changes at GCG sites can therefore be ruled out on three grounds.

**Table S1. Underlying genotypes, individuals, brain samples, datasets, and numbers of matches to AE4 and AE2 C versus T probes: primary data**

| Underlying genotype | | Individual | M/F | Disease status | SRA | Brain region | AE4 site: AE4=C (APOE4) C | AE4 site: AE4=T (APOE2+E3) T | AE2 site: AE2=C (APOE3+E4) C | AE2 site: AE2=T (APOE2) T | % C↔T |
|---|---|---|---|---|---|---|---|---|---|---|---|
| APOE4 | APOE4 | SD005/19 | M | AD | SRX17674439 | AMYG | 1191 | 59 | 1152 | 0 | **4.72** |
| | | | | | SRX17674440 | BA24 | 391 | 22 | 407 | 0 | **5.33** |
| | | | | | SRX17674438 | HPC | 491 | 49 | 542 | 1 | **9.07** |
| | | | | | SRX17674437 | HYPO | 804 | 83 | 990 | 0 | **9.36** |
| APOE4 | APOE3 | SD001/17 | F | AD | SRX17674435 | AMYG | 659 | 841 | 1131 | 28 | **2.42** |
| | | | | | SRX17674436 | BA24 | 752 | 960 | 1479 | 9 | **0.6** |
| | | | | | SRX17674467 | HPC | 794 | 1021 | 1374 | 30 | **2.14** |
| | | | | | SRX17674466 | HYPO | 485 | 533 | 740 | 21 | **2.76** |
| APOE4 | APOE3 | SD037/18 | M | AD | SRX17674443 | AMYG | 601 | 676 | 1187 | 1 | 0.08 |
| | | | | | SRX17674444 | BA24 | 483 | 591 | 851 | 0 | 0 |
| | | | | | SRX17674442 | HPC | 694 | 673 | 1307 | 2 | 0.15 |
| | | | | | SRX17674441 | HYPO | 773 | 841 | 1313 | 0 | 0 |
| APOE3 | APOE3 | SD030/18 | M | CTRL | SRX17674455 | AMYG | 3 | 1681 | 1452 | 0 | 0.18 |
| | | | | | SRX17674457 | BA24 | 0 | 1090 | 1039 | 0 | 0 |
| | | | | | SRX17674454 | HPC | 0 | 1178 | 946 | 0 | 0 |
| | | | | | SRX17674453 | HYPO | 12 | 1058 | 854 | 0 | **1.12** |
| APOE3 | APOE3 | SD035/15 | M | CTRL | SRX17674460 | AMYG | 2 | 852 | 788 | 1 | **0.23** |
| | | | | | SRX17674461 | BA24 | 2 | 1405 | 1105 | 1 | 0.14 |
| | | | | | SRX17674459 | HPC | 0 | 1202 | 807 | 0 | 0 |
| | | | | | SRX17674458 | HYPO | 0 | 1953 | 1574 | 0 | 0 |
| APOE3 | APOE3 | SD042/18 | F | CTRL | SRX17674452 | AMYG | 14 | 2543 | 1948 | 3 | **0.55** |
| | | | | | SRX17674451 | HPC | 26 | 2094 | 1650 | 1 | **1.23** |
| | | | | | SRX17674450 | HYPO | 29 | 1088 | 919 | 0 | **2.6** |
| APOE3 | APOE3 | SD014/17 | M | AD | SRX17674448 | AMYG | 23 | 410 | 344 | 0 | **5.31** |
| | | | | | SRX17674449 | BA24 | 5 | 245 | 247 | 0 | **2** |
| | | | | | SRX17674447 | HPC | 11 | 506 | 461 | 0 | **2.13** |
| | | | | | SRX17674446 | HYPO | 28 | 873 | 677 | 0 | **3.11** |
| APOE3 | APOE3 | SD032/17 | M | AD | SRX17674464 | AMYG | 1 | 3757 | 2789 | 5 | 0.03 |
| | | | | | SRX17674465 | BA24 | 0 | 880 | 718 | 0 | 0 |
| | | | | | SRX17674463 | HPC | 0 | 2619 | 1964 | 3 | 0 |
| | | | | | SRX17674462 | HYPO | 0 | 3371 | 2574 | 0 | 0 |
| APOE3 | APOE2 | SD025/19 | M | AD | SRX17674445 | AMYG | 2 | 3878 | 1401 | 1408 | 0.05 |
| | | | | | SRX17674456 | BA24 | 1 | 1302 | 670 | 546 | 0.08 |
| | | | | | SRX17674434 | HPC | 2 | 3012 | 1160 | 1167 | 0.07 |
| | | | | | SRX17674433 | HYPO | 2 | 1974 | 732 | 798 | 0.1 |

| | | | |
|---|---|---|---|
| M | Male | AMYG | Amygdala |
| F | Female | BA24 | Cingulate cortex |
| AD | Alzheimer's disease | HPC | Hippocampus |
| CTRL | Control | HYPO | Hypothalamus |

**Footnote:** The sequencing error rate in this series was ~0.05% per position, and therefore one expects ~1 change at a particular nucleotide in a dataset of ~1000–3000 *APOE* sequences. The frequency of pyrimidine to pyrimidine changes versus pyrimidine to purine changes is about one third of this (i.e., 0.33). We therefore introduced a cut-off of 10-fold above this (i.e., 5 specific replacements per dataset of 2000 *APOE* reads, and in multiple samples from the same individual), noting that a mean of 22 replacements were observed in the four key individuals, which is 66-fold (unnormalized) higher than the rate expected from sequencing errors.

**Table S2. Analysis of single-cell RNA sequencing data from brain microglia from control and Alzheimer's disease (AD) individuals**

| Individual | M/F | Age | Disease status | Region | Brain bank |
|---|---|---|---|---|---|
| 2018-018 | F | 82 | CTR | LPS | Netherlands Brain Bank |
| 2018-112 | F | 95 | CTR | LPS | Netherlands Brain Bank |
| iB6408-BA7 | M | 76 | CTR | LPS | Antwerp University Hospital |
| 2019-010 | F | 77 | MCI | LPS | Netherlands Brain Bank |
| 2018-120 | M | 72 | CTR+ | LPS | Netherlands Brain Bank |
| 2018-090 | M | 97 | CTR+ | LPS | Netherlands Brain Bank |
| 2018-021 | M | 92 | CTR+ | LPS | Netherlands Brain Bank |
| 2018-105 | M | 86 | CTR+ | LPS | Netherlands Brain Bank |
| 2018-135 | F | 81 | AD | LPS | Netherlands Brain Bank |
| 2019-030 | M | 77 | AD | LPS | Netherlands Brain Bank |
| iB6399-BA7 | F | 88 | AD | LPS | Antwerp University Hospital |
| 2018-064 | F | 103 | CTR+ | LPS | Netherlands Brain Bank |
| 2019-032 | M | 92 | AD | LPS | Netherlands Brain Bank |

**Table S3. List of single-cell microglia datasets analyzed in Figure 4 in the main text**

SRA numbers of single-cell microglia data are from (Alsema *et al.* 2020). The SRAs are presented in numeric order whereas the data in Figure 4 are sorted according to nucleotide abundance; not all SRAs are plotted in Figure 4 (extremely large numbers or absent readcounts have been removed to facilitate data plotting). The complete datasets can be found online at NCBI (https://www.ncbi.nlm.nih.gov/bioproject/?term=PRJNA611563).

| Individual | SRA | GEO ID |
|---|---|---|
| 2018-018 | SRX7878600 | GSM4403092 |
| 2018-018 | SRX7878601 | GSM4403093 |
| 2018-018 | SRX7878602 | GSM4403094 |
| 2018-018 | SRX7878603 | GSM4403095 |
| 2018-018 | SRX7878629 | GSM4403105 |
| 2018-018 | SRX7878630 | GSM4403106 |
| 2018-018 | SRX7878631 | GSM4403107 |
| 2018-018 | SRX7878640 | GSM4403108 |
| 2018-018 | SRX7878717 | GSM4403137 |
| 2018-021 | SRX7878604 | GSM4403096 |
| 2018-021 | SRX7878605 | GSM4403097 |
| 2018-021 | SRX7878606 | GSM4403098 |
| 2018-021 | SRX7878607 | GSM4403099 |
| 2018-021 | SRX7878644 | GSM4403112 |
| 2018-021 | SRX7878645 | GSM4403113 |
| 2018-021 | SRX7878646 | GSM4403114 |
| 2018-021 | SRX7878647 | GSM4403115 |
| 2018-021 | SRX7878747 | GSM4403207 |
| 2018-105 | SRX7878614 | GSM4403154 |
| 2018-105 | SRX7878618 | GSM4403158 |
| 2018-105 | SRX7878619 | GSM4403159 |
| 2018-105 | SRX7878620 | GSM4403160 |
| 2018-105 | SRX7878638 | GSM4403170 |
| 2018-105 | SRX7878672 | GSM4403180 |
| 2018-105 | SRX7878705 | GSM4403189 |
| 2018-105 | SRX7878708 | GSM4403192 |
| 2018-105 | SRX7878715 | GSM4403135 |
| 2018-105 | SRX7878720 | GSM4403196 |
| 2018-105 | SRX7878732 | GSM4403144 |
| 2018-105 | SRX7878818 | GSM4403234 |
| 2018-112 | SRX7878615 | GSM4403155 |
| 2018-112 | SRX7878639 | GSM4403171 |
| 2018-112 | SRX7878656 | GSM4403172 |
| 2018-112 | SRX7878673 | GSM4403181 |
| 2018-112 | SRX7878704 | GSM4403188 |
| 2018-112 | SRX7878707 | GSM4403191 |
| 2018-112 | SRX7878716 | GSM4403136 |
| 2018-112 | SRX7878722 | GSM4403198 |
| 2018-112 | SRX7878733 | GSM4403145 |
| 2018-112 | SRX7878750 | GSM4403210 |
| 2018-112 | SRX7878761 | GSM4403213 |
| 2018-112 | SRX7878763 | GSM4403215 |

| Individual | SRA | GEO ID |
|---|---|---|
| 2018-064 | SRX7878608 | GSM4403148 |
| 2018-064 | SRX7878624 | GSM4403100 |
| 2018-064 | SRX7878625 | GSM4403101 |
| 2018-064 | SRX7878626 | GSM4403102 |
| 2018-064 | SRX7878627 | GSM4403103 |
| 2018-064 | SRX7878636 | GSM4403168 |
| 2018-064 | SRX7878668 | GSM4403120 |
| 2018-064 | SRX7878669 | GSM4403121 |
| 2018-064 | SRX7878670 | GSM4403122 |
| 2018-064 | SRX7878671 | GSM4403123 |
| 2018-064 | SRX7878693 | GSM4403129 |
| 2018-064 | SRX7878718 | GSM4403138 |
| 2018-064 | SRX7878746 | GSM4403206 |
| 2018-090 | SRX7878609 | GSM4403149 |
| 2018-090 | SRX7878637 | GSM4403169 |
| 2018-090 | SRX7878662 | GSM4403178 |
| 2018-090 | SRX7878694 | GSM4403130 |
| 2018-090 | SRX7878706 | GSM4403190 |
| 2018-090 | SRX7878709 | GSM4403193 |
| 2018-090 | SRX7878719 | GSM4403139 |
| 2018-090 | SRX7878725 | GSM4403201 |
| 2018-090 | SRX7878727 | GSM4403203 |
| 2018-090 | SRX7878745 | GSM4403205 |
| 2018-090 | SRX7878751 | GSM4403211 |
| 2018-090 | SRX7878813 | GSM4403229 |
| 2018-120 | SRX7878655 | GSM4403243 |
| 2018-120 | SRX7878680 | GSM4403244 |
| 2018-120 | SRX7878698 | GSM4403254 |
| 2018-120 | SRX7878699 | GSM4403255 |
| 2018-120 | SRX7878721 | GSM4403197 |
| 2018-120 | SRX7878736 | GSM4403260 |
| 2018-120 | SRX7878752 | GSM4403268 |
| 2018-120 | SRX7878764 | GSM4403216 |
| 2018-120 | SRX7878777 | GSM4403221 |
| 2018-120 | SRX7878782 | GSM4403226 |
| 2018-120 | SRX7878817 | GSM4403233 |
| 2018-135 | SRX7878649 | GSM4403237 |
| 2018-135 | SRX7878681 | GSM4403245 |
| 2018-135 | SRX7878683 | GSM4403247 |
| 2018-135 | SRX7878696 | GSM4403252 |
| 2018-135 | SRX7878737 | GSM4403261 |

| | | |
|---|---|---|
| 2018-112 | SRX7878776 | GSM4403220 |
| 2018-112 | SRX7878819 | GSM4403235 |
| 2019-010 | SRX7878651 | GSM4403239 |
| 2019-010 | SRX7878654 | GSM4403242 |
| 2019-010 | SRX7878685 | GSM4403249 |
| 2019-010 | SRX7878687 | GSM4403251 |
| 2019-010 | SRX7878700 | GSM4403256 |
| 2019-010 | SRX7878753 | GSM4403269 |
| 2019-010 | SRX7878767 | GSM4403219 |
| 2019-010 | SRX7878775 | GSM4403283 |
| 2019-010 | SRX7878780 | GSM4403224 |
| 2019-010 | SRX7878781 | GSM4403225 |
| 2019-010 | SRX7878812 | GSM4403228 |
| 2019-010 | SRX7878816 | GSM4403232 |

| | | |
|---|---|---|
| 2018-135 | SRX7878740 | GSM4403264 |
| 2018-135 | SRX7878749 | GSM4403209 |
| 2018-135 | SRX7878754 | GSM4403270 |
| 2018-135 | SRX7878755 | GSM4403271 |
| 2018-135 | SRX7878760 | GSM4403212 |
| 2018-135 | SRX7878765 | GSM4403217 |
| 2018-135 | SRX7878778 | GSM4403222 |
| 2018-135 | SRX7878779 | GSM4403223 |
| 2018-135 | SRX7878783 | GSM4403227 |
| 2018-135 | SRX7878784 | GSM4403284 |
| 2018-135 | SRX7878815 | GSM4403231 |

**Table S4. Primary *APOE* sequence variation data for transgenic mice expressing either human *APOE4* or *APOE3*[a]**

| | | AE4 | | AE2 | |
|---|---|---|---|---|---|
| | | C | T/U | C | T/U |
| JOHNSON (KENTUCKY); LEE *et al.* | | | | | |
| SRA | GENOTYPE | | | | |
| SRX17356400 | E4 (C+C) | 5000 | 4 | 4999 | 5 |
| SRX17356401 | E4 | 5000 | 2 | 5000 | 2 |
| SRX17356403 | E4 | 5000 | 3 | 5000 | 2 |
| SRX17356404 | E4 | 3747 | 4 | 3869 | 2 |
| SRX17356405 | E4 | 5000 | 69 | 5000 | 2 |
| SRX17356408 | E4 | 5000 | 12 | 5000 | 7 |
| SRX17356410 | E4 | 4624 | 21 | 4650 | 7 |
| SRX17356413 | E4 | 5000 | 10 | 4800 | 1 |
| SRX17356414 | E4 | 4508 | 19 | 4333 | 1 |
| SRX17356415 | E4 | 3417 | 18 | 3244 | 6 |
| SRX17356418 | E4 | 4237 | 25 | 4503 | 3 |
| SRX17356399 | E3 (T+C) | 1 | 3265 | 4035 | 7 |
| SRX17356402 | E3 | 17 | 3837 | 4149 | 0 |
| SRX17356406 | E3 | 20 | 5000 | 5000 | 0 |
| SRX17356407 | E3 | 4 | 3975 | 4016 | 2 |
| SRX17356409 | E3 | 17 | 5000 | 5000 | 0 |
| SRX17356411 | E3 | 5 | 5000 | 5000 | 3 |
| SRX17356412 | E3 | 68 | 4273 | 3994 | 2 |
| SRX17356416 | E3 | 30 | 4391 | 4180 | 0 |
| SRX17356417 | E3 | 2 | 3475 | 3856 | 3 |
| TAVAZOIE (ROCKEFELLER); MILLET *et al.* | | | | | |
| SRX21236909 | E4 (C+C) | 4678 | 8 | 4098 | 1 |
| SRX21236907 | E4 | 4398 | 33 | 3663 | 1 |
| SRX21236905 | E4 | 2433 | 0 | 2039 | 5 |
| SRX21236906 | E4 | 4958 | 19 | 4393 | 3 |
| SRX21236908 | E4 | 4415 | 16 | 3821 | 2 |
| SRX21236903 | E3 (T+C) | 4 | 5000 | 5000 | 23 |
| SRX21236902 | E3 | 2 | 5000 | 4992 | 8 |
| SRX21236900 | E3 | 2 | 3638 | 3043 | 23 |
| SRX21236904 | E3 | 2 | 5000 | 5000 | 9 |
| SRX21236901 | E3 | 2 | 5000 | 5000 | 10 |

MEAN SEQUENCE ERROR RATE 0.0447%

SD 0.0603%

ERROR RATE + 2xSD 0.165%

[a]Values give the number of perfect matches using probes selective for different variants at the AE4 and AE2 sites. Source RNA-seq data from (Lee *et al.* 2023) and (Millet *et al.* 2024). 5000 is the maximum number of matches permitted by BLAST at NCBI, but from the published sizes of the databases it can be calculated that the number of expected matches would at most be only slightly above (<5%).

**Figure S1.** Specificity of sequence changes at the AE2 site in individual human brain microglia, and variation in per-cell expression levels.

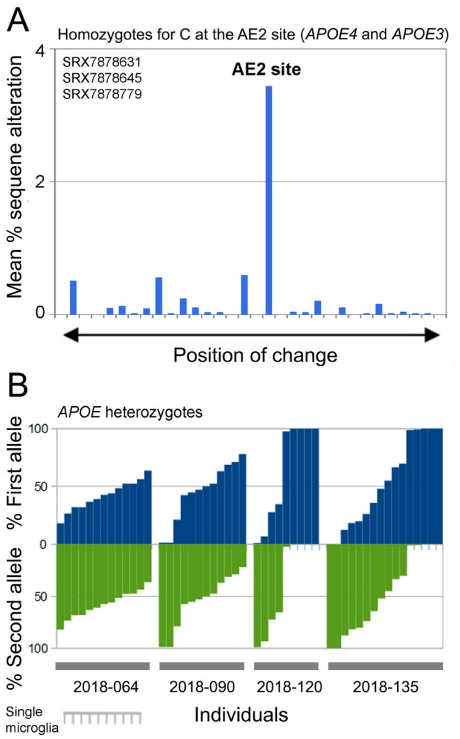

**Legend:** (A) The three datasets in Figure 4 with evidence of C to T changes at the AE2 site (red in Figure 4B) were analyzed for the locations of all sequence replacements across the AE2 region as described in Figure 2B. This demonstrates that the majority of sequence changes affect the specific AE2 nucleotide and are thus not sequencing errors. (B) Variation in per-cell expression levels. For the four APOE heterozygous individuals in Figure 4 (right), data were replotted as the percentage of mRNA expressed from each allele, and sorted left to right in ascending order, which revealed that the per-cell expression of a given allele varies from 0% to 100%, indicative of monoallelic expression (MAE)/random allelic expression (RAE). Because the mean number of readcounts (excluding zeros) was ~300 (see Figure 4), the finding is unlikely to reflect sparse coverage.

**Figure S2.** Studies on *APOE* ribotype in transgenic mice expressing human *APOE4*.

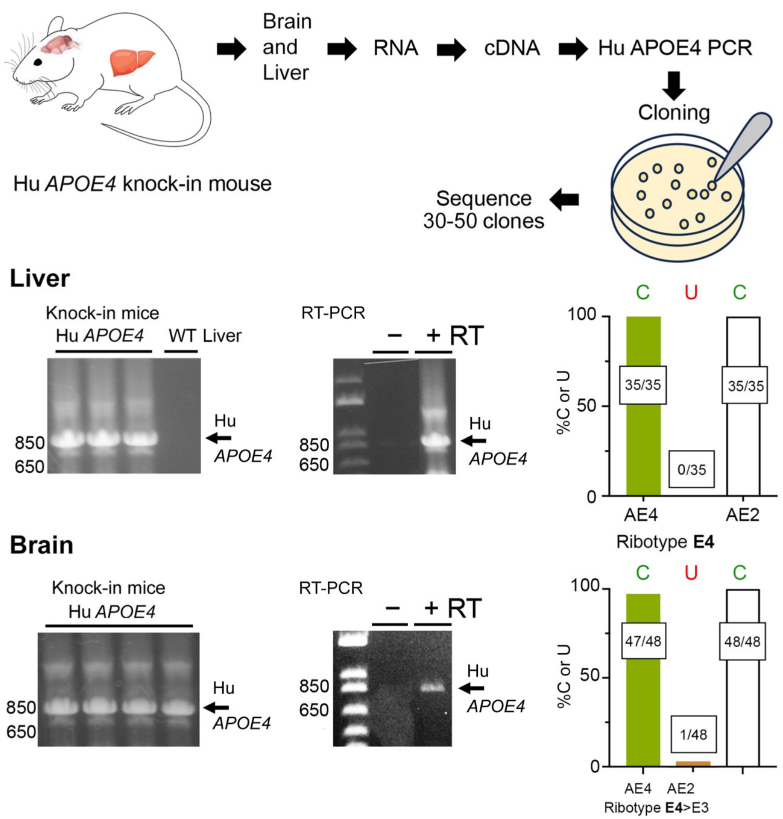

**Legend:** Low-level sequence change at the AE4 site in brain but not liver of transgenic mice harboring human (Hu) *APOE4*. (Top panel) Schematic: liver and brain RNA from *APOE4* knock-in mice was converted to cDNA and the region comprising AE4 and AE2 was PCR amplified, cloned, and sequenced. (Left panels) Confirmation of PCR amplification of *APOE* (gel electrophoresis), no PCR product was obtained from wild-type (WT) mice. (Center panels) No PCR product was obtained when reverse transcripase (RT) was omitted, confirming that only mRNA is being analyzed. (Right panels) No changes were seen at the AE4 site in mouse liver, but one of 48 clones (~2%) from brain contained a C→T (i.e., C→U) change at this site. From the sequence error rate this change would only be expected once in ~50 experiments, suggestive of a low level of sequence replacement, which has now been confirmed by further data (main text). No sequence changes were observed at AE2 or at any other sites.

**Figure S3.** Non-genomic transcripts in transgenic mice expressing human *APOE4*.

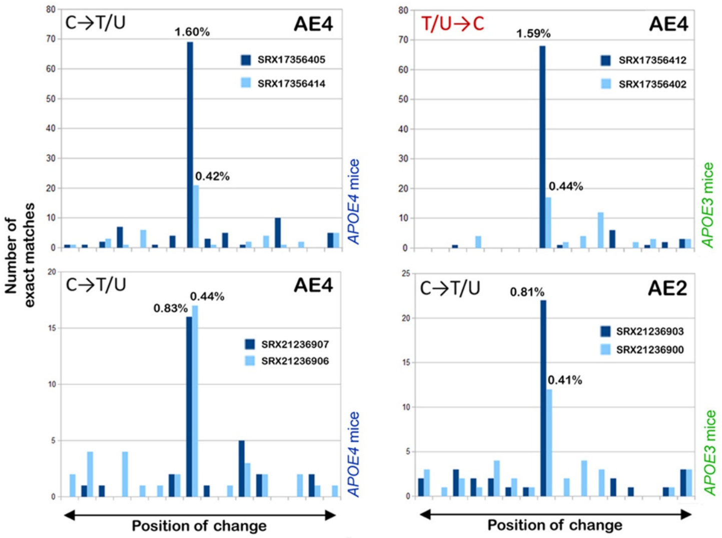

**Legend.** Sequence replacements in brains of mice harboring human *APOE*. Sequence changes in *APOE* mRNA across the AE4 and AE2 sites (analysis as in Figure 2) from brain tissue (four samples) or cortical microglia (four samples) of transgenic mice harboring human *APOE4* or *E3* (genotype shown on the right); source data from Johnson and colleagues (Lee *et al.* 2023) and Tavazoie and colleagues (Millet *et al.* 2024) filed online at NCBI. The *y* axis shows the absolute number of readcounts (rather than percentages) to emphasize that the results are based on relatively large numbers of reads; figures over the central bars give the percent replacement at the AE4 or AE2 sites. Values exceeding 0.2% are considered to be statistically significant (sequence error rate 0.045% + 2×SD = 0.165%; $P$ = <0.05); no other sites reproducibly exceeded this threshold. Abbreviation: SD, standard deviation.

**Figure S4.** Unexplained variation in *APOE* genotype and ribotype in human liver-derived cells

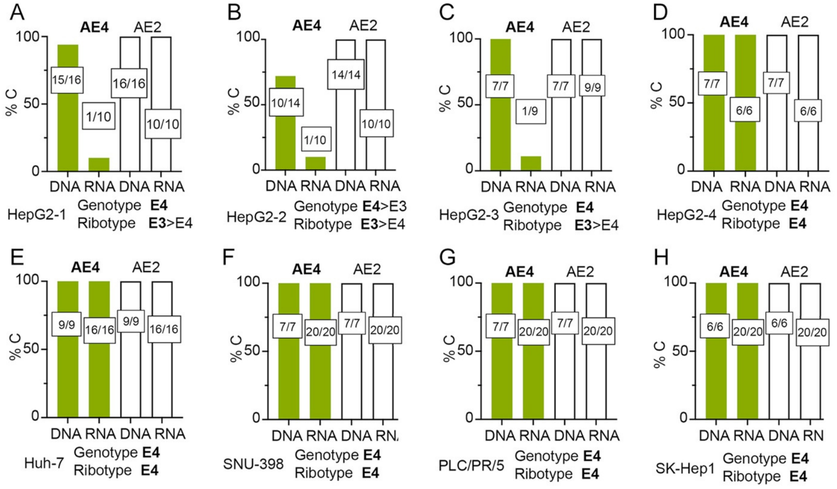

**Legend:** Human cell lines express a diversity of *APOE* alleles which can influence their utility (Schaffer *et al.* 2014). We therefore validated the *APOE* genotype and ribotype from human Huh-7 and HepG2 liver-derived cells which were reported to be *APOE3/3* (Schaffer *et al.* 2014). The figure plots the % C at the AE4 and AE2 sites, all deviations were C→U (i.e., T in cDNA). Four different clones of HepG2 cells from the same source (ATCC, Methods) that have been maintained over more than 20 years were analyzed, along with several other human liver-derived cell lines. (A–D) Among the four different HepG2 clonal lines, and contrary to our expectations, all exhibited predominantly (clones 1–3) or (D) exclusively an *APOE4* genotype (clone 4). Among the clones exhibiting a polymorphic *APOE* genotype, all three (clones 1–3) exhibited a C→U change at the AE4 site and a predominantly (~90%) APOE3 ribotype, that was unrelated to passage number or date of acquisition. The fourth HepG2 clone exhibited both *APOE4* genotype and ribotype. (E) Our isolate of Huh-7 was revealed to be exclusively *APOE4*/*APOE4*, and all the other liver-derived cell lines (F–H) examined exhibited both *APOE4* genotype and ribotype.

These findings suggest that *APOE* is unpredictably polymorphic at the AE4 site in some clones of HepG2 cells, with an unexplained discrepancy between genotype (*APOE4*) and ribotype (*APOE3*) that diverges from a report that HepG2 cells are exclusively of the *APOE3*/*APOE3* genotype (Schaffer *et al.* 2014). Nevertheless, other data confirm that HepG2 cells across the world are predominantly of the *APOE3*/*APOE3* genotype (supplementary data below).

| **Supplementary data:** Many HepG2 cells are *APOE3* homogygotes BLAST searching of RNA-seq data with allele-specific probes | AE4 | | AE2 | | Genotype |
|---|---|---|---|---|---|
| | C | T | C | T | |
| **North Carolina State University** | | | | | |
| SRX28167007 | 2 | >500 | >500 | 2 | *APOE3*/*APOE3* |
| SRX28167006 | 1 | >500 | >500 | 3 | *APOE3*/*APOE3* |
| **U1103 INSERM, Clermont-Ferrand, France** | | | | | |
| SRX26353534 | 0 | 16 | 3 | 0 | *APOE3*/*APOE3* |
| SRX26353527 | 0 | 146 | 16 | 0 | *APOE3*/*APOE3* |
| **Zhejiang University** | | | | | |
| SRX28586134 | 6 | >500 | >500 | 32 | *APOE3*/*APOE3* |
| SRX28586133 | 7 | >500 | >500 | 34 | *APOE3*/*APOE3* |
| **Catholic University Liver Research Center, South Korea** | | | | | |
| SRX24720382 | 1 | >500 | >500 | 0 | *APOE3*/*APOE3* |
| SRX24720381 | 1 | >500 | >500 | 1 | *APOE3*/*APOE3* |

The unexpected deviation of the genoypes of our established cell lines, all from ATCC, but which have been maintained in culture for many years, leads us to suspect that *in vitro* selection could have led to changes at the *APOE* locus. Although our cultures are routinely maintained with appropriate antibiotics, microbiome analysis revealed that they are not microbiologically sterile (not presented), an issue that has been recognized for many years (Fogh 1973).

**Figure S5.** Screening of the entire *APOE* and *APP* transcripts for single-nucleotide changes.

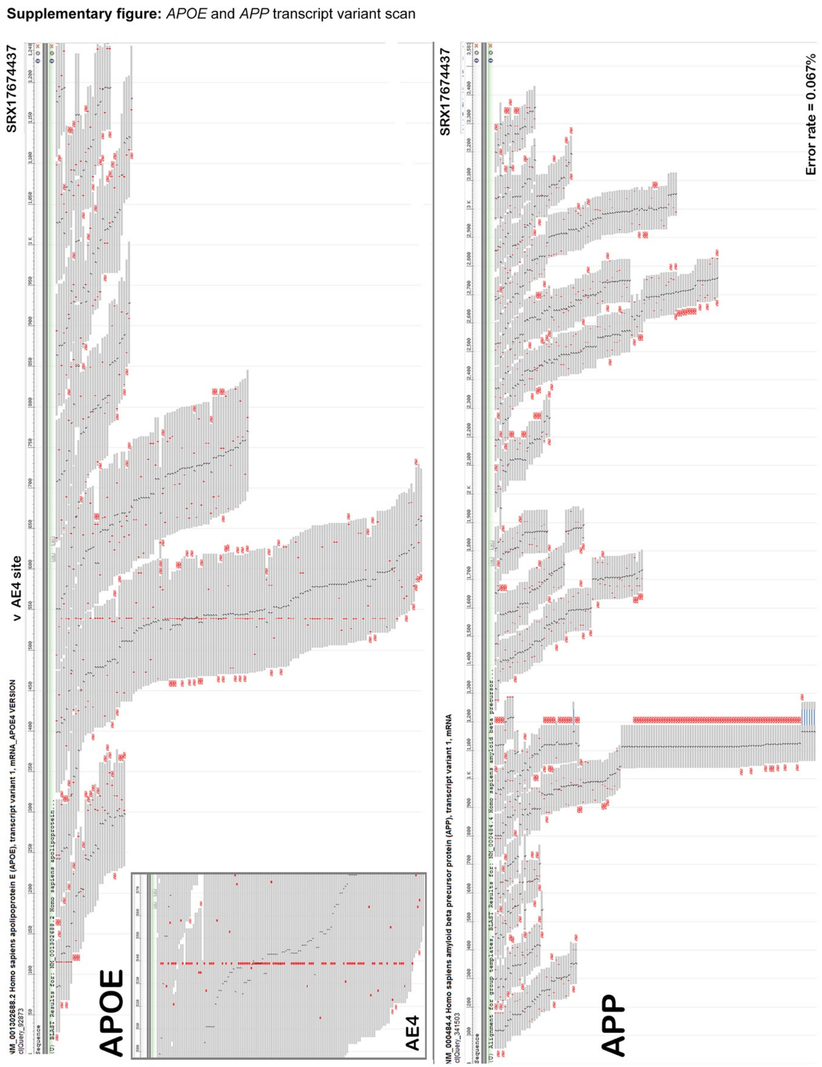

**Legend:** Location of all single-nucleotide changes (red squares; red symbols are discontinuities caused by differential splicing) in full-length *APOE* and *APP* mRNAs (SRAs are given at the top of each panel). In this analysis only one site is systematically altered in *APOE*, and no such site is present in *APP*.

**Figure S6.** Locations CpG nucleotides in *APOE* cDNA

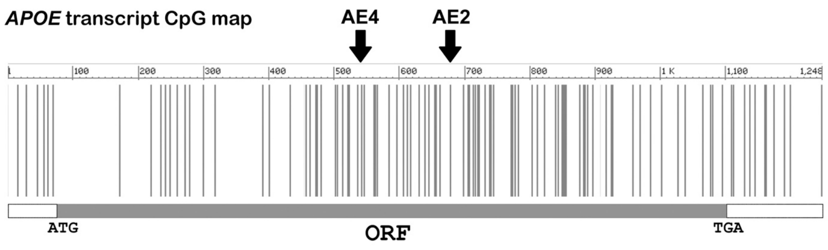

**Legend:** Methylation of cytosines (mC) in DNA makes them potentially prone to spontaneous hydrolytic deamination, generating thymine residues. Both inferred editing sites are in CpG nucleotides, where the ancestral form *APOE4* is C and the human-specific versions E3 and E2 represent stepwise replacement of CpG by TpG. Although *APOE* is CpG-methylated in brain (Yu *et al.* 2013), and there are potential mechanisms to convert mC to T, only the AE4 and AE2 sites are altered. This high specificity argues against CpG-based mechanisms, and that DNA methylation does not explain the observed C→T/U transitions (nor the T/U→C changes).

**Figure S7.** Analysis of potential antisense transcription of human *APOE.*

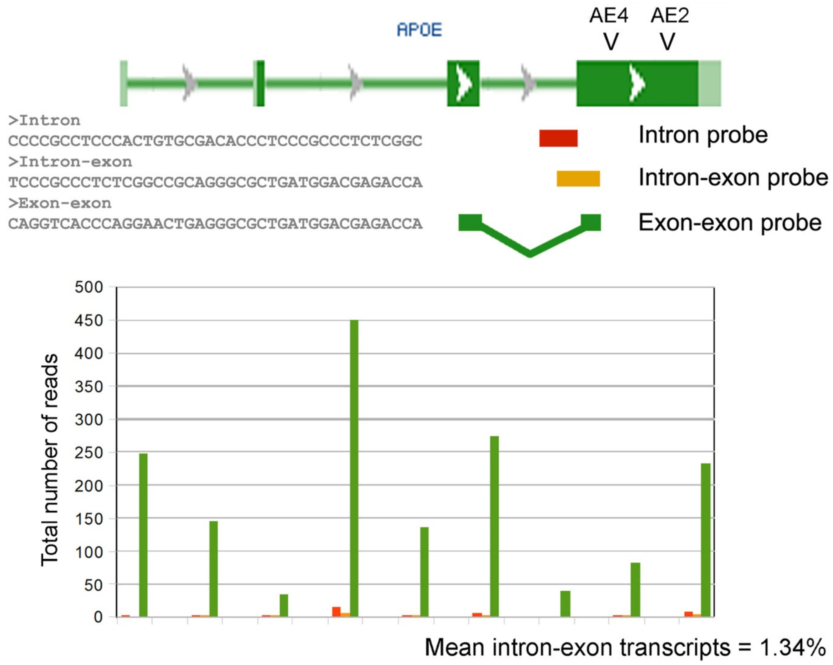

**Legend.** RNA-seq datasets (e.g., SRA entries) generally do not report the strand of origin. To determine whether there is a significant level of antisense transcription for APOE we reasoned that antisense transcripts would not respect splice junctions, and therefore analyzed the first RNA-seq dataset in each case from the nine individuals listed in Table S1 using probes that overlapped the intron-exon boundary adjacent to the exon containing AE4 and AE2. Note that BLAST finds an identical number of matches using sense or antisense probes. This analysis revealed that only a low level of transcripts cross the intron-exon boundary. This could be due to either unspliced mRNA or to antisense transcription, but sets an upper limit to the extent of any antisense transcription.

We also studied potential antisense microRNAs. We were unable to confirm the reported (Pencheva *et al.* 2012) homology between miR-1908 or miR-199a and *APOE* mRNA. BLAST searching revealed that their level of expression in brain is 1.39% of *APOE*; it is therefore unlikely that they make a major contribution to RNA editing.

**Figure S8.** APOBEC enzymes expressed in brain.

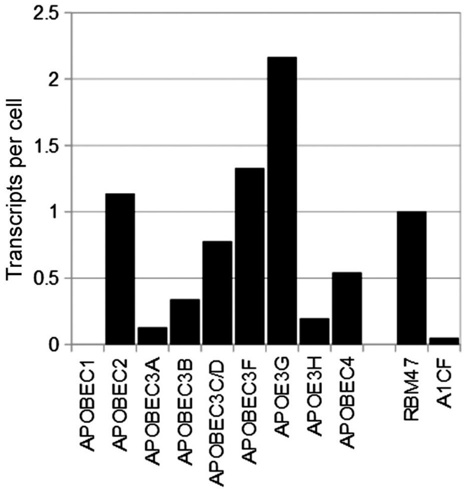

**Legend.** Mean mRNA expression levels of APOBEC family enzymes and related factors in human brain samples ($N$ = 35). Data were normalized to levels of well-expressed housekeeping genes (*PGK1*, *HMGCR*, *NSE1*; ca 50 transcripts per cell (Hu *et al.* 2022)) to determine the absolute number of *APOBEC* transcripts per cell. *APOBEC3C* and *APOBEC3D* have been pooled because of their extensive sequence co-identity. APOBEC3G is known to be expressed and functionally active in brain neurons (Hill *et al.* 2006; Wang *et al.* 2009), but APOBEC1 cannot be excluded as a candidate because of remarkable disparities between levels of APOBEC enzymes detected by RNA expression, mass spectrometry, and immunohistochemistry, as illustrated by studies on APOBEC3A (https://www.proteinatlas.org/ENSG00000128383-APOBEC3A/tissue).
